# Th2 cell extracellular vesicles promote eosinophil survival through the cytokine cargo IL-3 and prolong airway eosinophilia

**DOI:** 10.1101/2024.07.23.600647

**Authors:** Kaitlyn E. Bunn, Brenna G. Giese-Byrne, Heather H. Pua

## Abstract

**Background:** Extracellular vesicles (EVs) mediate intercellular communication during immune responses. EVs are abundant in respiratory biofluids, and the composition of EVs in the lung changes during inflammation.

**Objective:** We aimed to quantify the contribution of T cells to airway EVs in allergic lung inflammation and ascertain their function during a type 2 inflammatory response.

**Methods:** Genetic membrane tagging was combined with single vesicle flow cytometry to quantify T cell EVs in the airways of mice challenged with ovalbumin or house dust mite. EVs were purified from T helper type 2 (Th2) cell cultures and their functions on eosinophils assessed by flow cytometry and RNA sequencing. Th2 cell EVs were instilled into the lungs of mice to determine effects on lung eosinophilia. Finally, the function of an EV protein cargo was tested using inhibitors and blocking antibodies.

**Results:** T cell EVs are increased in the airways of mice with induced allergic inflammation. EVs secreted by Th2 cells inhibit apoptosis and induce activating pathways in eosinophils *in vitro.* This effect depends on re-stimulation through the T cell receptor. Th2 cell EVs prolong eosinophilia *in vivo* during allergic airway inflammation. Th2 cell EVs carry a potent form of the cytokine IL-3 on their surfaces, which inhibits apoptosis by activating Jak1/2-dependent pro-survival programs in eosinophils.

**Conclusion:** Th2 cell EVs promote eosinophil survival and prolong eosinophilia during allergic airway inflammation. This function depends on the EV cargo IL-3, supporting a role for EVs as vehicles of cytokine-based communication in lung inflammation.

**Key Messages:**

- T cells secrete extracellular vesicles in the airway during allergic lung inflammation.
- Th2 cell extracellular vesicles inhibit eosinophil apoptosis and prolong airway eosinophilia during allergic lung inflammation.
- IL-3 carried on Th2 cell EVs is a functional cargo, supporting a role for cytokine-carrying EVs as drivers of type 2 inflammation.

**Capsule summary:** This study supports that T cell extracellular vesicles may be important drivers of eosinophilic inflammation through the cytokine cargo IL-3, offering new insights into pro-inflammatory signaling in the allergic lung of patients with asthma.

## Introduction

Asthma is a multicellular disease driven by complex communication networks. In allergic asthma, T helper type 2 (Th2) cells can recruit and activate other cells of the type 2 inflammatory response, including eosinophils, mast cells, and IgE-producing B cells.^1^ Increases in secreted cytokines, chemokines, and bioactive lipids promote this inflammation and are targets for biologic therapies.^2,3^ Extracellular vesicles (EVs) have recently emerged as a new class of secreted factors present in lung inflammation. EVs are small, lipid bilayer delimited particles secreted by cells that participate in cell-cell communication through their bioactive cargos.^4^ Respiratory biofluids contain abundant EVs, with concentrations ranging from 10^8^ to 10^10^ EVs/ml.^5–7^ Studies in mice and humans have shown that EVs from epithelial cells, endothelial cells, and immune cells are present in respiratory fluids and that the composition of airway EVs changes with inflammation.^8,9^ In particular, EVs originating from immune cells are increased in respiratory biofluids collected from patients with chronic rhinosinusitis, bronchopulmonary dysplasia, and asthma.^10–12^

In previous work, we found that immune cell EVs and their microRNA cargos are increased in a mouse model of allergic airway inflammation.^13^ Immune cell EVs not only mark biofluids during inflammation, but also potently mediate inter-immune cell communication and function in diverse immunological processes including antigen presentation,^14–16^ allorecognition,^17,18^ anti-tumor immunity,^19–21^ immune cell chemotaxis,^22–24^ and immunosuppression.^25,26^ EVs have also been shown to participate in allergic immune responses through antigen presentation to allergen-specific T cells and the promotion of Th2 cell polarization.^27–29^ Thus, EVs likely constitute an important axis of communication in asthma, though the individual functions of immune cell EVs are incompletely understood.

Like other immune cells, T cells secrete EVs. These EVs have been shown to carry various cargos involved in T cell immune functions, including cytokines, chemokines, microRNAs, tRNA fragments, DNA, integrins, and the T cell receptor (TCR) complex.^30–35^ Functionally, T cell EVs can mediate communication between T cells and other immune cells through delivery of suppressive cargos like microRNAs, cytokines, Fas ligand, and APO2 ligand as well as activating cargos like IFNγ to target cells.^36–39^ Despite the known importance of T cells in driving allergic airway inflammation, whether T cells contribute to airway EVs during type 2 inflammation in the lung and the functional roles T cell EVs serve are relatively unknown. In this study, we investigated the role of T cell EVs in promoting eosinophilia in allergic airway inflammation. Using genetic membrane tagging and functional studies both *in vitro* and *in vivo*, we determined that T cells secrete EVs in the allergic lung and that EVs secreted by Th2 cells re-activated through the T cell receptor can promote eosinophil survival and eosinophilia in the lung through the cytokine cargo IL-3.

## Methods

### Animals

All animal studies were performed after obtaining approval from the Vanderbilt Institutional Animal Care and Use Committee.

### *In vivo* allergic airway studies

For ovalbumin (OVA), mice were sensitized intraperitoneally with 50 µg OVA mixed with Alum, challenged with 50 µg OVA by oropharyngeal aspiration on days 7, 8, and 9 following sensitization, and euthanized on day 10. For house dust mite (HDM), mice were challenged with 40 µg HDM extract by oropharyngeal aspiration three times per week for three weeks and then euthanized the day following the last challenge. For papain, mice were challenged with 3.5 µg papain by oropharyngeal aspiration and euthanized 24 to 120 hours following challenge. Bronchoalveolar lavage fluid (BALF) and lungs were collected, processed, and analyzed by flow cytometry.

### Purification of EVs from primary Th2 cell cultures

CD4^+^ T cells were isolated from mouse spleens and lymph node single cell suspensions by positive bead selection. Isolated cells were polarized to Th2 cells by stimulation with anti-CD3, anti-CD28, IL-4, and anti-interferon gamma for 3 days. On day 3, cells were removed from anti-CD3/anti-CD28 stimulation and rested in media with IL-4 and IL-2.

On day 5, cell culture media was centrifuged at 250xg for 10 minutes to pellet cells. The supernatant was collected and centrifuged at 2,000xg for 30 minutes to pellet large cellular debris and was used for purification of rested Th2 cell EVs. The cells were then stimulated with anti-CD3, anti-CD28, and IL-2 for 6 hours, and media was subjected to 250xg and 2,000xg differential centrifugation and was used for isolation of activated Th2 cell EVs. EVs were isolated from cell culture supernatant by size exclusion chromatography (SEC) using Izon qEV columns per manufacturer instructions.

Fractions were concentrated using 100 kDa ultrafiltration. EVs were characterized by nanoparticle tracking analysis, transmission electron microscopy, and single vesicle flow cytometry.

### *In vitro* eosinophil viability studies

Eosinophils were differentiated from mouse bone marrow by culturing bone marrow cells with FLT-3 ligand and SCF on days 0-3 and IL-5 on days 4-12. Eosinophils were characterized by microscopy, flow cytometry, and functional studies. On culture day 12, mature eosinophils were incubated with 5×10^9^ EVs/ml of rested and activated Th2 cell EVs for 6 or 24 hours, and viability was assessed by membrane permeability, caspase 3 activity, and annexin V binding. For ruxolitinib studies, eosinophils were treated with 1.6 µM ruxolitinib or DMSO vehicle control for 30 minutes at 37C prior to the addition of EVs. For IL-3 blocking studies, EVs were incubated with anti-IL-3 or isotype control antibody for 30 minutes at 37C prior to eosinophil treatment.

### Bulk RNA-sequencing

Eosinophils treated with EVs from activated Th2 cell culture or parallelly processed cell culture media for 24 hours were homogenized, and total RNA was isolated and submitted to Vanderbilt Technologies for Advanced Genomics Core for sequencing on the NovaSeq 6000. Data were processed using the lab’s RNA-seq pipeline (https://github.com/parkdj1/RNASeq-Pipeline). Primary data were deposited in the Gene Expression Omnibus (https://www.ncbi.nlm.nih.gov/geo/). Raw and processed data can be accessed using the accession number GSE253180 with the reviewer token ojexoeygplqbrgv. Downstream analysis included differential expression using the R package DESeq2 (https://www.r-project.org/).

### Statistical Analysis

Data were expressed as mean values plus or minus standard errors of the mean or as median and interquartile range depending on the normality of the data. Statistical significance was determined through tests denoted in the figure legends performed by GraphPad Prism 10 software (GraphPad Software, La Jolla, CA). P-value of < 0.05 was considered statistically significant.

## Supplementary methods

Additional methodologic details for T cell and eosinophil culture, EV purification, EV characterization (nanoparticle tracking analysis, transmission electron microscopy, single vesicle flow cytometry), mouse allergic airway models, *in vivo* administration of EVs, cell viability assays, IL-3 detection and blocking, and RNA-sequencing are provided within the supplement.

## Results

### T cell EVs are increased in the allergic lung

To test whether T cell EVs are produced in the allergic lung, we utilized a strategy previously developed in the lab to trace the cellular origins of EVs *in vivo*.^13^ We crossed *mTmG* mice^40^ with *Lck-Cre* mice^41^ to switch the expression of membrane-targeted Tomato (mTomato) for membrane-targeted GFP (mGFP) in T cells expressing Cre recombinase (**Fig. 1A**). As expected, GFP uniquely labeled T cells in these mice (**Supplemental Fig. 1A**). By single vesicle flow cytometry, we detected mGFP^+^ EVs from *in vitro* cultured *Lck-Cre mTmG* CD4^+^ T cells, and these mGFP^+^ EVs could be degraded by the detergent Triton X-100 but not proteinase K (**Fig. 1B, C)**. GFP in EVs purified by SEC also co-fractionated with the EV marker ALIX and the T cell membrane markers CD11a and CD71, without contaminating subcellular membranes (Lamin B1-nucleus, Tomm20-mitochondria) (**Fig. 1D**).

**Figure 1.**
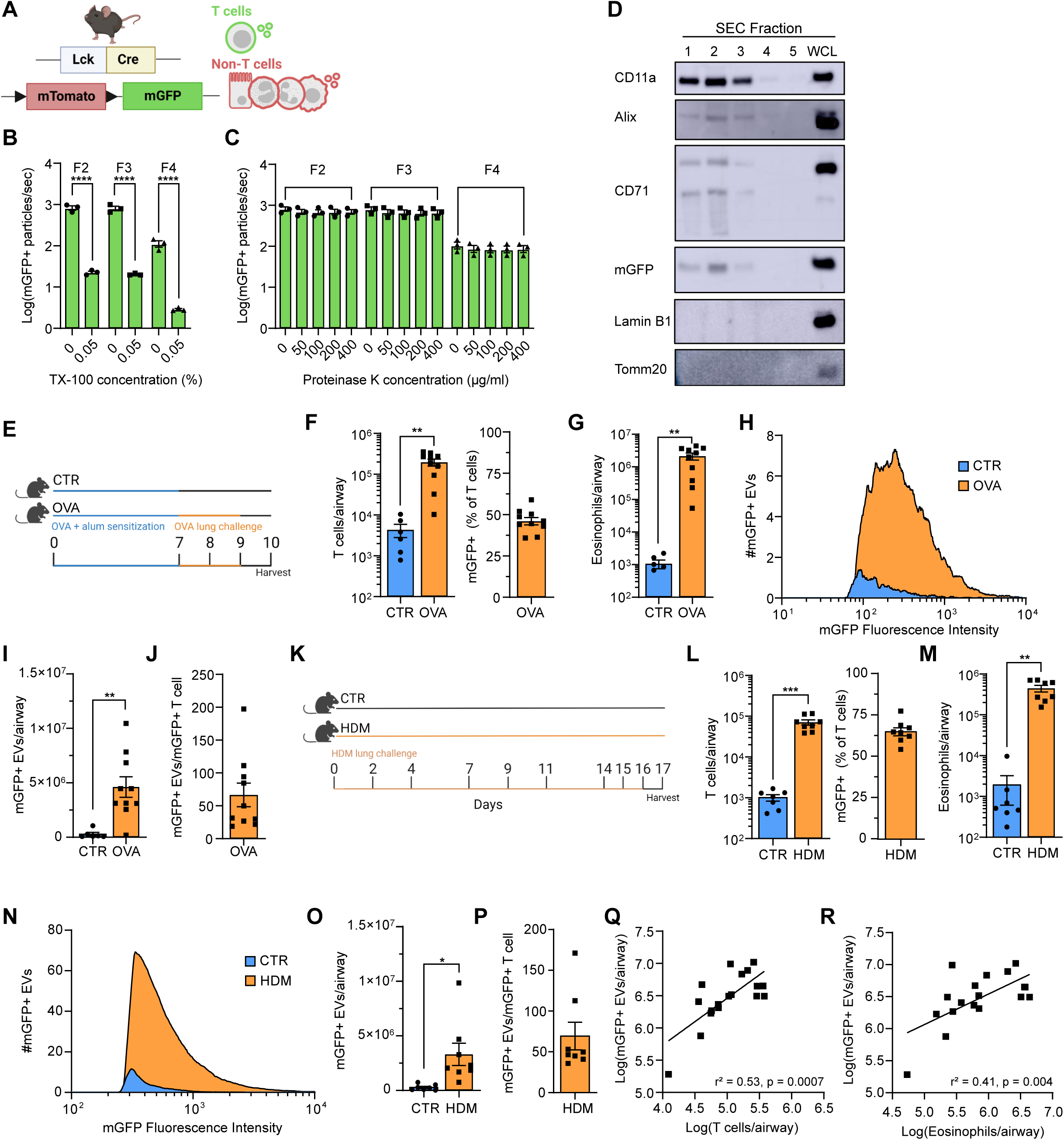
T cell EVs are increased in the allergic lung. **(A)** Graphical depiction of the *Lck-Cre mTmG* mouse model, in which T cell membranes and T cell EVs are labeled with membrane-targeted GFP (mGFP) and all other cells are labeled with membrane-targeted Tomato (mTomato). **(B, C)** Quantification of mGFP^+^ particles detected per second in *Lck-Cre mTmG* T cell culture media by single vesicle flow cytometry following **(B)** Triton X-100 treatment and **(C)** proteinase K treatment of varying concentrations. N = 3, 2-way ANOVA with Sidak test for multiple comparisons. **(D)** Western blot of lysates obtained by SEC fractionation of *Lck-Cre mTmG* Th2 cell culture supernatant at 6 hrs of culture. Equal loading volume between fractions. WCL = whole cell lysate. **(E)** Graphical depiction of ovalbumin (OVA) allergic airway induction model, CTR = control (vehicle-challenged). **(F)** Quantification of the number of T cells (left) and the percentage of mGFP^+^ T cells in the airways of CTR and OVA mice following allergic airway induction. **(G)** Quantification of the number of eosinophils in the airways of CTR and OVA mice following allergic airway induction. **(H)** Averaged histograms depicting the number of EVs corresponding to mGFP fluorescence intensity in CTR and OVA mice detected by single vesicle flow cytometry in BALF. **(I)** Quantification of mGFP^+^ particles in the airways of CTR and OVA mice following allergic airway induction. **(J)** Quantification of the number of mGFP^+^ particles per mGFP^+^ T cell in the airways of CTR and OVA mice following allergic airway induction. **(K)** Graphical depiction of house dust mite (HDM) allergic airway induction model, CTR = control (vehicle-challenged). **(L)** Quantification of the number of T cells (left) and the percentage of mGFP^+^ T cells in the airways of CTR and HDM mice following allergic airway induction. **(M)** Quantification of the number of eosinophils in the airways of CTR and HDM mice following allergic airway induction. **(N)** Averaged histograms depicting the number of EVs corresponding to mGFP fluorescence intensity in CTR and HDM mice detected by single vesicle flow cytometry in BALF. **(O)** Quantification of mGFP^+^ particles in the airways of CTR and HDM mice following allergic airway induction. **(P)** Quantification of the number of mGFP^+^ particles per mGFP^+^ T cell in the airways of CTR and HDM mice following allergic airway induction. **(Q, R)** Quantification of the correlation between mGFP^+^ particles and T cells **(Q)** or eosinophils **(R)** in the airways of allergic airway mice (OVA and HDM experiments combined). **(F-J)** n = 6 CTR, n = 10 OVA, 2 independent experiments, two-tailed t test. **(L-P)** n = 7 CTR, n = 8 HDM, 2 independent experiments, two-tailed t test. **(Q, R)** n = 10 OVA, 8 HDM, 4 independent experiments, simple linear regression. All error bars represent SEM, ** p < 0.01, *** p < 0.001. By flow cytometry, T cells were defined as TCRβ^+^ and CD4^+^ or CD8^+^, and eosinophils were defined as Siglec F^+^ CD11b^+^ SSC^high^.

Having confirmed that EVs secreted by *Lck-Cre mTmG* T cells are labeled with GFP, we next induced allergic airway inflammation in *Lck-Cre mTmG* mice by ovalbumin (OVA) or house dust mite (HDM) sensitization and challenge (**Fig. 1E, K**). Cellular flow cytometry on BALF showed a significant increase in airway T cells and eosinophils in OVA- and HDM-challenged mice, indicating the development of allergic, type 2 inflammation (**Fig. 1F, G, L, M**), with 50% of airway T cells labeled with mGFP (**Fig. 1F, L**). To quantify mGFP labeled EVs in the airway, we performed single vesicle flow cytometry on BALF. We detected a mean of 4.62×10^6^ and 3.31×10^6^ mGFP^+^ EVs/airway in OVA- and HDM-challenged mice, respectively, which was significantly higher than in control mice (**Fig. 1H, I, N, O**). The linear range of detection of mGFP^+^ EVs from BALF was established by serial dilution studies, and the limit of detection was defined by running filtered PBS blanks (**Supplemental Fig. 1B**). Loss of mGFP after Triton X-100 treatment supported that mGFP in BALF of *Lck-Cre mTmG* mice was carried by membrane-bound EVs (**Supplemental Fig. 1C**). When the number of T cell EVs were compared relative to the number of T cells in the airway, there were on average 67 and 69 GFP^+^ EVs/GFP^+^ T cell in OVA- and HDM-challenged mice, respectively (**Fig. 1J, P**).

The number of GFP^+^ T cell EVs in the BALF of allergic airway mice positively correlated with both the number of T cells and the number of eosinophils in the BALF (**Fig. 1Q, R**). Together, these data show that T cells secrete EVs in the allergic airway during active lung inflammation.

### Th2 cell EVs promote eosinophil survival *in vitro*

Given the critical role of Th2 cells in driving allergic airway inflammation, we next investigated EV secretion from *in vitro* polarized Th2 cells that were either re-stimulated through the T cell receptor (“activated”) following polarization or not (“rested”) (**Fig. 2A**). To avoid contamination from serum-derived EVs, T cells were polarized in serum-free media, and Th2 polarization was confirmed by cytokine secretion (**Supplemental Fig. 2A, B**). By single vesicle flow cytometry in *Lck-Cre mTmG* mice, mGFP^+^ T cell EV secretion did not differ between rested and activated Th2 cells up until 24 hours of culture (**Fig. 2B, Supplemental Fig. 2C, D**), at which point activated Th2 cells showed increased cell death (**Fig. 2C, Supplemental Fig. 3A**). Increased EV secretion by activated Th2 cells at 24 hours may be due to increased debris associated with cell death; thus, we used a 6-hour activation timepoint for our studies. Nanoparticle tracking analysis confirmed there was no difference in quantity of EVs secreted by rested and activated Th2 cells at 6 hours of culture (**Fig. 2D**). By nanoparticle tracking analysis and transmission electron microscopy, there were also no size or morphological differences between rested and activated Th2 cell EVs (**Supplemental Fig. 3B-D**).

**Figure 2.**
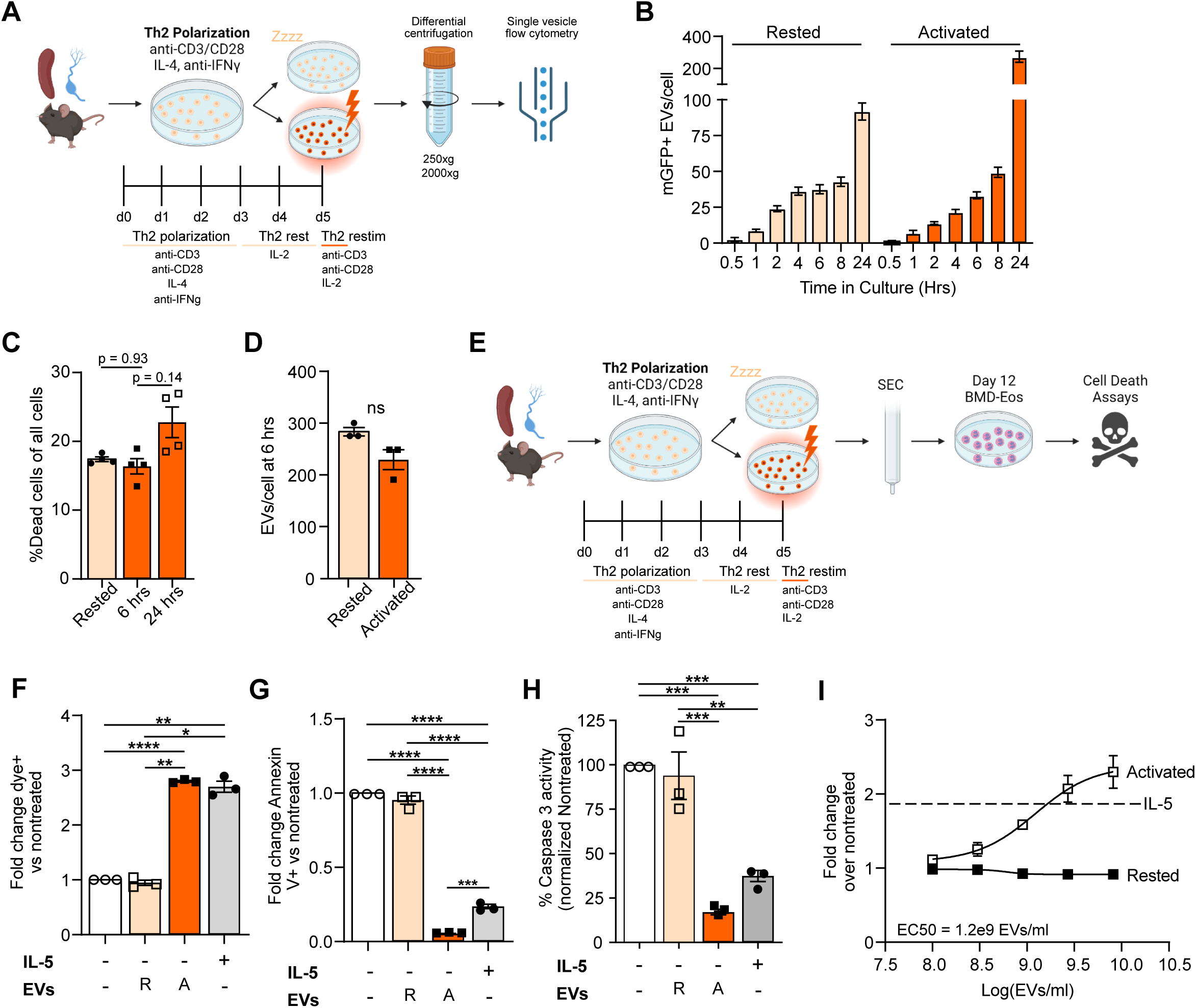
Th2 cells promote eosinophil survival *in vitro*. **(A)** Graphical depiction of EV purification from rested and activated Th2 cell culture media and subsequent single vesicle flow cytometry. **(B)** Quantification of EV production over time by rested and activated Th2 cells by single vesicle flow cytometry. n = 3 for 0.5-8 hrs, n = 2 for 24 hrs. **(C)** Quantification of T cell death at 6 and 24 hrs following activation. n = 4, one-way ANOVA (vs. rested) with Bonferroni’s correction for multiple testing. **(D)** Quantification of EV production by rested and activated Th2 cells at 6 hrs of culture by NTA. n = 3, two-tailed t test. **(E)** Graphical depiction of EV purification from rested and activated Th2 cell culture media and day 12 bone marrow-derived eosinophil cell death assays following Th2 cell EV treatment. **(F)** Quantification of eosinophil survival by dye permeability assay at 24 hrs post-Th2 cell EV treatment. **(G)** Quantification of Annexin-V^+^, permeability dye^-^ eosinophils following 6 hrs of Th2 cell EV treatment. **(H)** Quantification of caspase-3 activity in eosinophils following 6 hrs of Th2 cell EV treatment. **(I)** Quantification of eosinophil survival following 24 hrs of treatment with increasing doses of Th2 cell EVs, n = 3, log[agonist] vs response with variable response (4 parameters). For **(F-H)**, n = 3 independent experiments each with 3 eosinophil biologic replicates and one pooled sample of T cell EVs (5,000 EVs/cell). One-way ANOVA with Tukey’s multiple comparisons test. All error bars represent SEM, * p < 0.05, ** p < 0.01, *** p < 0.001, **** p < 0.0001.

Because a hallmark of allergic airway inflammation is lung eosinophilia, we investigated the effects of Th2 cell EVs on eosinophils. Mouse eosinophils differentiated from bone marrow (**Supplemental Fig. 4A-D**) were treated with EVs purified by SEC from rested and activated Th2 cell culture media, and viability was assessed with a membrane permeable dye (**Fig. 2E**). We treated eosinophils with 5×10^9^ EVs/ml, 2x our estimated physiological concentration of T cell EVs in airway lining fluid during allergic inflammation (**Supplemental Fig. 4E**). Treatment with rested EVs did not change survival from the nontreated condition; however, treatment with activated Th2 cell EVs increased eosinophil survival 2.5x, equivalent to the survival benefit conferred by exogenous dosing of the canonical pro-survival cytokine IL-5 (**Fig. 2F**). Activated Th2 cell EVs promoted eosinophil survival through the inhibition of apoptosis, with decreased levels of Annexin V binding and decreased caspase 3 activity in treated eosinophils (**Fig. 2G, H**). This effect on viability was dose dependent, with a half maximal effective concentration (EC50) of 1.2×10^9^ EVs/ml (**Fig. 2I**). No effects of rested Th2 cell EVs were seen after treating eosinophils with increasing doses, suggesting a qualitative rather than quantitative difference in EVs secreted following Th2 cell activation through the TCR.

### Th2 cell EVs prolong eosinophilia *in vivo*

After establishing that activated Th2 cell EVs increase eosinophil viability *in vitro*, we hypothesized that activated Th2 cell EVs would prolong eosinophilia during allergic airway inflammation *in vivo*. To minimize the contribution of endogenous T cell derived EVs, we employed the papain model of allergic airway inflammation, in which damage to the airway epithelium caused by papain’s protease activity leads to the recruitment of eosinophils in absence of an endogenous T cell response.^42^ In our papain model, eosinophil numbers peaked at 48 hours following papain challenge and largely resolved by 72 hours (**Fig. 3A, B**). We found that administration of 2×10^9^ activated Th2 cell EVs at 48 hours after papain administration resulted in significantly increased SiglecF^+^CD11b^+^SSC^high^ eosinophils at 72 hours in the airways when compared to mice treated with parallelly processed unconditioned media as a vehicle control (**Fig. 3C, D**). Interestingly, there was also a trend towards increased tissue eosinophils in the lung in EV treated mice, whereas there was no difference in the number of eosinophils in the blood vessels in the lungs (**Fig. 3E, F**). Total airway and lung tissue F4/80^+^CD11c^+^ macrophages and TCRβ^+^CD4^+^ T cells did not change with EV treatment, indicating a selective effect on eosinophils (**Fig. 3G-J**). Together these data demonstrate that EVs produced from TCR-activated Th2 cells can prolong eosinophil-rich inflammation in the lung.

**Figure 3.**
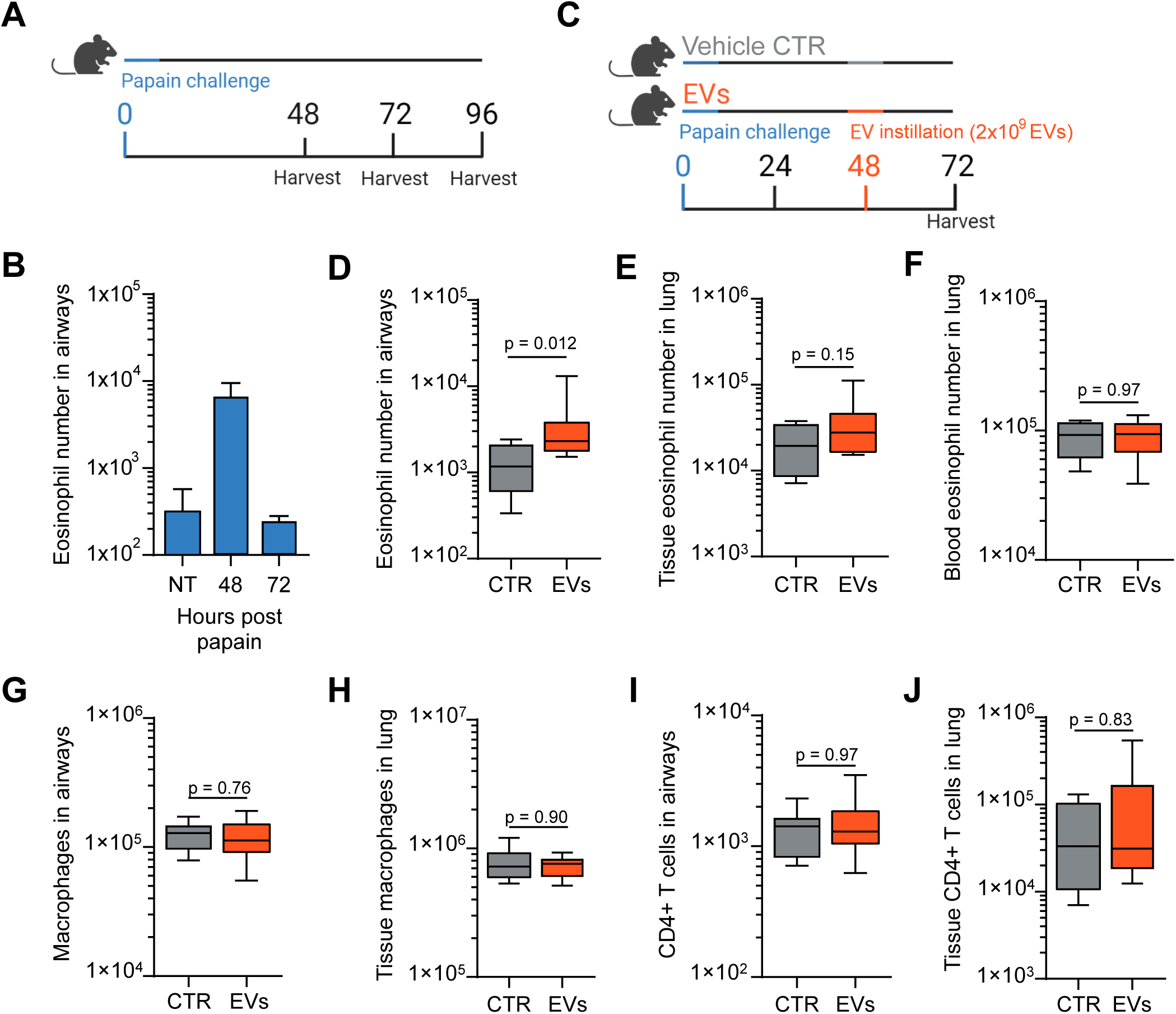
Th2 cell EVs prolong allergic eosinophilia *in vivo*. **(A)** Graphical depiction of papain administration and BALF collection along a time course to assess eosinophil kinetics after aspiration of a single dose of 3.5 µg papain. **(B)** Quantification of BALF eosinophils in mice not treated with papain and at 48 and 72 hours following papain treatment. n = 4. **(C)** Graphical depiction of papain administration and activated Th2 cell EV or vehicle control challenge. **(D-F)** Quantification of **(D)** BALF eosinophils, **(E)** lung tissue eosinophils, and **(F)** blood eosinophils in the lungs of vehicle and Th2 cell EV treated mice. **(G, H)** Quantification of macrophages in the **(G)** BALF and **(H)** lungs of vehicle and Th2 cell EV treated mice. **(I, J)** Quantification of CD4^+^ T cells in the **(I)** BALF and **(J)** lungs of vehicle and EV treated mice. For **(D-J)**, data are represented as median and IQR, n = 8 vehicle, 10 EV treated, two-tailed Mann-Whitney U test. By flow cytometry, eosinophils were defined as SiglecF^+^ CD11b^+^ SSC^hi^, macrophages as F4/80^+^ CD11c^+^, and CD4^+^ T cells as TCRβ^+^ CD4^+^.

### Th2 cell EVs induce survival and activation pathways in eosinophils

To determine how Th2 cell EVs affect eosinophil viability, we assessed eosinophil gene expression by bulk RNA sequencing following treatment of cultured eosinophils with activated Th2 cell EVs or parallelly processed unconditioned media as a vehicle control. Principal component analysis showed that 96% of variance was due to EV treatment (**Fig. 4A**). In EV- vs vehicle-treated eosinophils, 369 genes were significantly (adjusted p-value < 0.01) upregulated (Log_2_FC ≥ 1), and 667 genes were significantly downregulated (Log_2_FC ≤ -1) (**Fig. 4B, Supplemental Tables 1, 2**).

**Figure 4.**
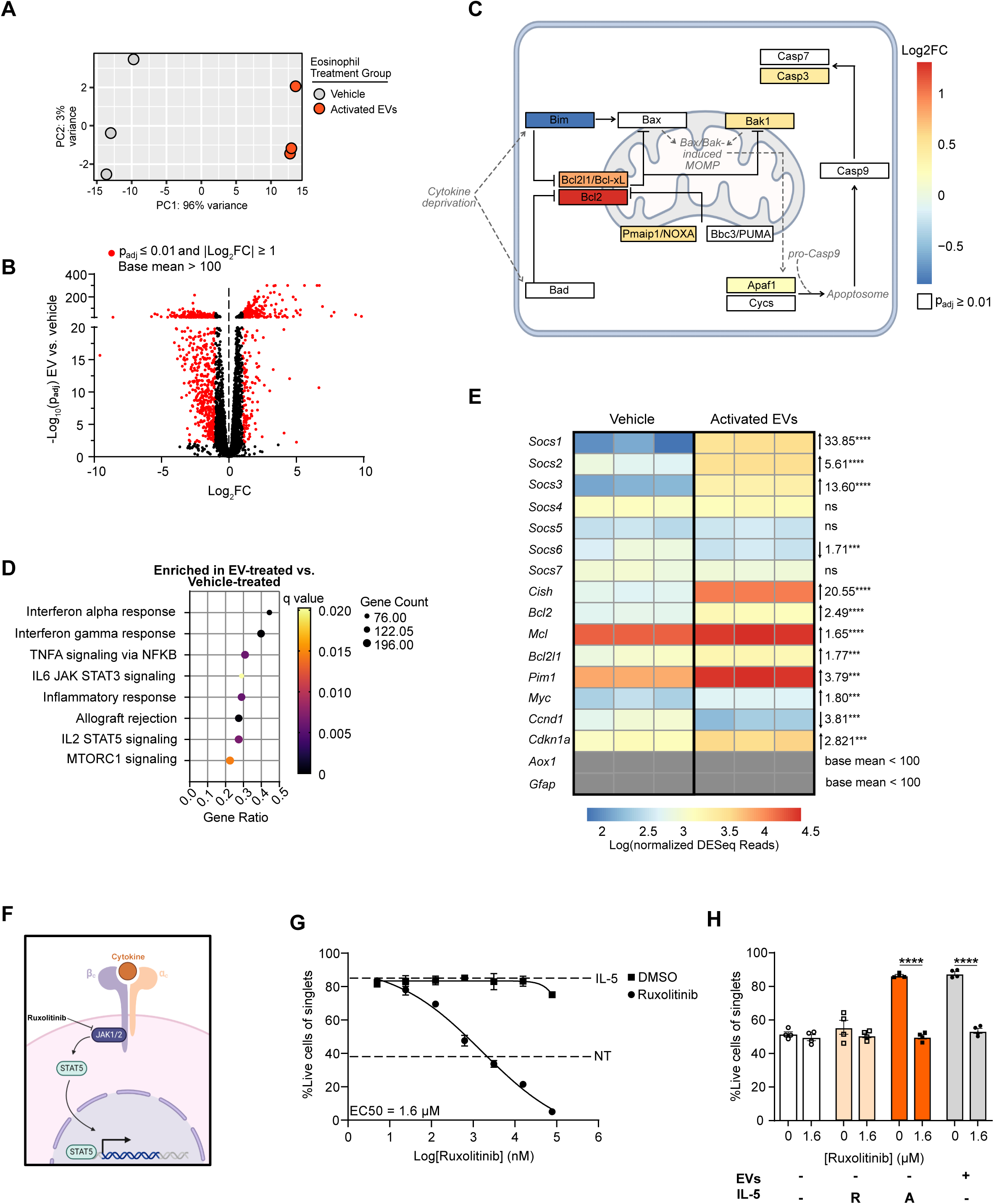
Th2 cell EVs induce survival and activation pathways in eosinophils. **(A)** Principal component analysis of gene expression differences between vehicle treated eosinophils (n = 3) and activated Th2 cell EV treated eosinophils (n = 3). **(B)** Volcano plot depicting differential gene expression between vehicle treated and activated Th2 cell EV treated eosinophils. Genes with adjusted p value ≤ 0.01 and |Log_2_FC| ≥1 are shown in red. **(C)** Intrinsic apoptosis diagram depicting KEGG intrinsic apoptosis pathway genes and their relationships with one another and corresponding Log_2_ fold change in gene expression between activated Th2 cell EV treated and vehicle treated eosinophils. Genes that are upregulated in EV treated eosinophils are shown in boxes with warm colors, genes that are downregulated in EV treated eosinophils are shown with cool colors, and genes that are not differentially expressed are shown in white. **(D)** Gene set enrichment analysis of activated Th2 cell EV treated vs vehicle treated eosinophils. Shown are gene sets with q-value ≤ 0.05. **(E)** Heatmap depicting KEGG Jak/Stat signaling pathway genes and corresponding normalized DESeq reads for those genes. Statistical analysis was performed using the R package DESeq2, and all p-values are adjusted for multiple testing. **(F)** Diagram depicting Jak1/2 signaling downstream of a common beta chain family cytokine binding to its receptor. **(G)** Quantification of eosinophil survival following 24 hrs treatment with increasing doses of ruxolitinib or vehicle (DMSO) and 10 ng/mL IL-5, n = 3, log[agonist] vs response with variable response (4 parameters). **(H)** Quantification of eosinophil survival following 24 hrs treatment with rested or activated Th2 cell EVs or IL-5 and 1.6 µM ruxolitinib, n = 4, 2-way ANOVA with Sidak’s correction for multiple comparisons. All error bars represent SEM, *** p < 0.001, **** p < 0.0001.

Consistent with protection of eosinophils from apoptosis with activated Th2 cell EV treatment (**Fig. 2G, H**), the intrinsic anti-apoptotic genes *Bcl2* and *Bcl2l1* (*Bcl-xL*) showed increased expression, and the intrinsic pro-apoptotic gene *Bcl2l11* (*Bim*) showed decreased expression in EV- vs vehicle-treated eosinophils (**Fig. 4C**). These genes are known to be regulated downstream of eosinophil survival pathways, including cytokine signaling, Fas/FasL signaling, CD40/CD40L signaling, TNF-α/fibronectin signaling, and Siglec-F crosslinking.^43–45^ We observed fewer significant changes in the extrinsic apoptosis pathway (**Supplemental Fig. 5A**).

Gene set enrichment analysis identified enrichment of gene sets involved in Jak/Stat signaling in EV- vs vehicle-treated eosinophils (**Fig. 4D**). Consistent with this observation, we saw upregulation of many genes downstream of Jak/Stat signaling, including the pro-survival factors *Mcl-1*, *Pim-1*, and *c-Myc* and the negative regulators of cytokine signaling *Socs1/2/3* (**Fig. 4E**). To test whether Jak/Stat signaling was required for eosinophil survival in response to activated Th2 cell EVs, we used the small molecule Jak1/2 inhibitor ruxolitinib (**Fig. 4F**), which has an EC50 of 1.6 µM in our eosinophil cultures (**Fig. 4G**). Eosinophils treated with 1.6 µM ruxolitinib prior to treatment with activated Th2 cell EVs showed complete abrogation of the survival benefit conferred by activated Th2 cell EVs (**Fig. 4H, Supplemental Fig. 5B**), indicating that activated Th2 cell EVs mediate eosinophil survival through Jak1/2 signaling.

Additional gene sets enriched in EV- vs vehicle-treated eosinophils included inflammatory/immunity-related sets – TNFα, interferon, inflammatory response, and allograft rejection (**Fig. 4D**). Several metabolism pathways were also regulated by EV treatment, with mTORC1 signaling genes increased and glycolysis genes decreased in EV treated cells (**Fig. 4D, Supplemental Fig. 5C**). Together, these data support that coordinated regulation of apoptosis in addition to inflammatory and metabolism pathways may allow eosinophils to both survive and participate in inflammatory responses after EV-mediated signaling from activated Th2 cells.

### IL-3 cargo on Th2 cell EVs promotes eosinophil survival

Jak/Stat signaling can be induced in eosinophils by the common beta chain cytokines, GM-CSF, IL-5, and IL-3, which are produced by activated Th2 cells and inhibit eosinophil apoptosis.^46–48^ Although common beta chain cytokines program many overlapping gene expression changes in eosinophils, IL-3 controls a distinct subset of genes different from GM-CSF and IL-5.^49^ Our RNA-sequencing data showed that activated Th2 cell EV treated eosinophils upregulate a subset of these genes uniquely induced by IL-3 but not IL-5 and GM-CSF (**Fig. 5A**). This included *Socs1* and *Socs3*, negative regulators of IL-3-induced cytokine signaling,^50^ *Upp1*, a uridine salvaging enzyme,^51^ *Ikzf4*, a STAT5-interacting transcription factor,^52^ *Pim2*, an anti-apoptotic gene,^53,54^ *CD69*, an activation marker,^55,56^ and *Xbp1*, a transcription factor indispensable for eosinophil differentiation (**Fig. 5B**).^57^ Consistent with this IL-3 gene expression signature, we found by ELISA that activated Th2 cell EVs carried an average of 6.9 pg IL-3 per 10^9^ EVs (min = 1.73, max = 21.3, SEM = 4.8), while IL-3 could not be detected on rested EVs (**Fig. 5C**). Lysed and unlysed EVs yielded equivalent IL-3 levels, consistent with IL-3 being carried largely on the surface of EVs (**Supplemental Fig. 6A, B**). IL-3 was enriched in fractions containing the highest EV particle counts by nanoparticle tracking analysis (**Fig. 5D**). We did not detect IL-3 in EV-containing SEC fractions collected from media conditioned by rested Th2 cells, consistent with production of IL-3 in T cells only after activation through the TCR (**Supplemental Fig. 6C**).^58^

**Figure 5.**
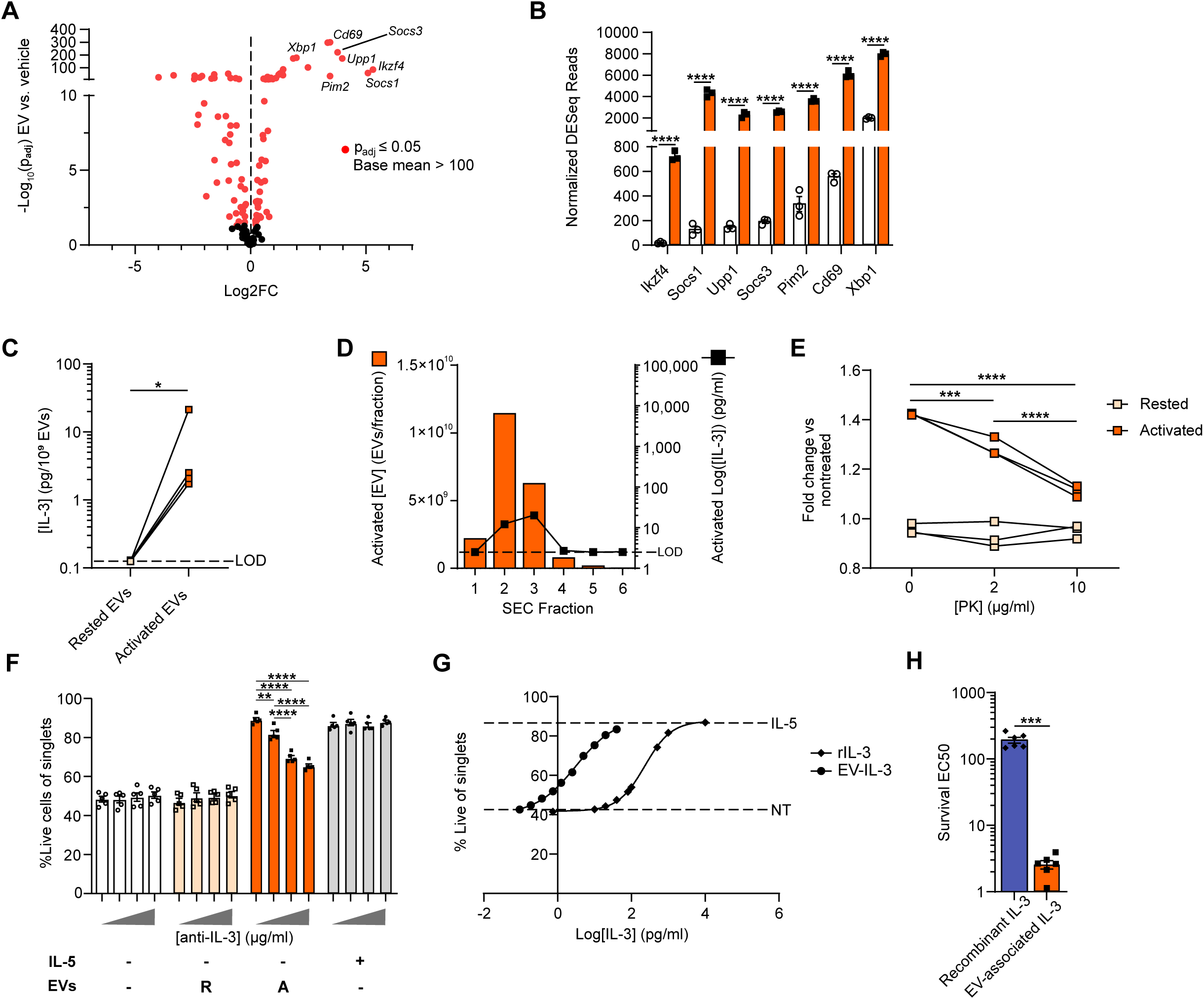
IL-3 cargo on Th2 cell EVs promotes eosinophil survival. **(A)** Volcano plot depicting differential gene expression between vehicle treated and activated Th2 cell EV treated eosinophils. Genes depicted are genes distinctly regulated by IL-3 per Nelson et al.^49^ Genes with adjusted p-value ≤ 0.05 are shown in red. **(B)** Normalized DESeq read counts for genes of interest upregulated in activated Th2 cell EV treated eosinophils vs vehicle treated eosinophils. Statistical analysis was performed using the R package DESeq2, and all p-values are adjusted for multiple testing. **(C)** Quantification of IL-3 present on SEC-purified rested and activated Th2 cell EVs measured by ELISA. n = 4, paired ratio two-tailed t test. The limit of detection (LOD) is defined by the manufacturer as 2.5 pg. **(D)** Quantification of EV number by NTA (left y-axis) and IL-3 concentration by ELISA (right y-axis) in SEC fractions 1-6 collected from activated Th2 cell culture media. The LOD is defined by the manufacturer as 0.125 pg/mL. **(E)** Quantification of eosinophil survival following 24 hrs treatment with Th2 cell EVs subjected to prior proteinase K treatment compared to non-treated eosinophils, n = 3, one-way ANOVA with Tukey’s multiple comparisons test. **(F)** Quantification of eosinophil survival following 24 hrs treatment with rested or activated EVs or vehicle (PBS) pre-incubated with increasing concentrations of anti-IL-3 (0, 1, 10, and 100 µg/mL). n = 5, 2-way ANOVA with Tukey’s correction for multiple comparisons, 2 independent experiments **(G)** Quantification of eosinophil survival following 24 hrs treatment with increasing doses of activated Th2 cell EV-associated IL-3 (0.09375 pg/mL, 0.1875 pg/mL, 0.375 pg/mL, 0.75 pg/mL, 1.25 pg/mL, 2.5 pg/mL, 5 pg/mL, 10 pg/mL, 20 pg/mL, and 40 pg/mL) or soluble recombinant IL-3 (0.75 pg/mL, 1.25 pg/mL, 2.5 pg/mL, 5 pg/mL, 10 pg/mL, 20 pg/mL, 40 pg/mL, 80 pg/mL, 100 pg/mL, 500 pg/mL, 1000 pg/mL, and 10,000 pg/mL), n = 3, log[agonist] vs response with variable response (4 parameters), representative of 2 independent experiments. **(H)** Comparison of eosinophil survival EC50 of soluble recombinant IL-3 or activated Th2 cell EV-associated IL-3, n = 6, 2 independent experiments, paired two-tailed t test. All error bars represent SEM, * p < 0.05, ** p < 0.01, *** p < 0.001, **** p < 0.0001.

Next, we tested whether EV-associated IL-3 is required for EVs to promote the survival of eosinophils. Following surface shaving of EVs by proteinase K, activated Th2 cell EVs did not support eosinophil survival, indicating that a surface protein cargo was required for EV function (**Fig. 5E, supplemental Fig. 6D-G**). Furthermore, IL-3 antibody blocking significantly decreased the survival of eosinophils treated with activated Th2 cell EVs in a dose-dependent manner (**Fig. 5F**). An isotype control antibody did not affect eosinophil survival (**Supplemental Fig. 6H**). Because EV-associated IL-3 promoted eosinophil survival at the low concentration of ∼35 pg/ml (equivalent to 6.9 pg IL-3 per 10^9^ EVs) (**Fig. 5C**), we hypothesized that EV-associated cytokine produced by activated Th2 cells was highly potent. Comparing dose response curves for EV-associated and recombinant IL-3 showed that the curve of EV-associated IL-3 was shifted dramatically to the left, indicating higher potency of EV-associated IL-3 than free recombinant cytokine (**Fig. 5G**). Activated Th2 cell EVs conferred a significant survival benefit at an IL-3 concentration of 0.19 pg/ml, while recombinant IL-3 did not significantly affect survival until a concentration of 20 pg/ml (**Fig. 5G**). The survival EC50 of EV-associated IL-3 was 3.1 pg/ml (95% CI 2.3 to 4.8 pg/ml), while the survival EC50 of recombinant IL-3 was 230 pg/ml (95% CI 210 pg/ml to 253 pg/ml), a 74x difference (**Fig. 5H**). These data suggest that activated Th2 cell EVs carry a potent form of IL-3 on their surfaces that promotes eosinophil survival.

## Discussion

Although therapies aimed at reducing eosinophil number are approved for clinical use in type 2 high asthma,^59^ eosinophils can remain in the lung and retain an activated phenotype in treated patients, suggesting the presence of untargeted drivers of disease-causing eosinophilia.^60^ Therefore, new mechanistic insights are needed to understand additional signals in type 2 inflammation to continue to improve the treatment of allergic airway disease. EVs have recently been identified as a form of cell-to-cell communication important for tissue inflammation.^61^ Both in individuals with asthma and in mouse models of allergic airway inflammation, we and others have shown that EV origins, numbers, and cargos are changed in respiratory biofluids.^12,13,62–64^ However, whether and how EVs may communicate signals in the airway remain poorly understood.

In this study, we used membrane tracing strategies to identify that T cell EVs are increased >10-fold in OVA and HDM models of allergic airway inflammation. Furthermore, our results support that activated Th2 cell EVs are sufficient to promote eosinophil survival and prolong eosinophilia in the lung through paracrine signaling. Our data also raise the possibility that EVs administered into the airspace may affect eosinophil numbers beyond the airway lumen in the lung parenchyma, raising important questions about the biodistribution of EVs relevant for both understanding pathophysiology and therapeutic applications in lung inflammation. These findings together with our observations that Th2 cell EVs are sufficient to induce robust pro-survival, pro-inflammatory, and metabolic gene expression programs in eosinophils support the hypothesis that Th2 cell EVs participate as a critical signaling axis and a part of the unique local microenvironment in type 2 lung inflammation.

This work also specifically highlights a role for T cell EVs in driving allergic lung inflammation by orchestrating other effector immune cells. Here we found that although activation of Th2 cells through the TCR does not change the basic physical characteristics (i.e. size, number, morphology) of secreted EVs, activated Th2 cell EVs are uniquely capable of promoting eosinophil survival, whereas EVs secreted by rested Th2 cells are not. These data suggest that polarized effector T cells in the tissues may need to encounter antigen to license EVs to promote inflammation. Future work is needed to test the function of EVs from other polarized effector cell populations (i.e. Th1 and Th17) in airway inflammation, how cues from the environment may change EV secretion in allergen-driven airway inflammation, and whether the function of T cell EVs changes in patients with asthma.

Finally, secreted cytokines have emerged as EV cargos, including T cell EVs.^30,37,39^ In this study, we found that activated Th2 cell EVs carry the common beta chain cytokine IL-3 which was required for their function in promoting eosinophil survival. EV-associated IL-3 was remarkably potent, with an effective concentration nearly 2 logs lower than recombinant IL-3. This raises important questions about whether EV-associated IL-3, even at low concentrations, may be a driver of lung inflammation in asthma, as IL-3 receptor is upregulated on lung eosinophils in patients with asthma.^65^ Since EVs may allow cytokines to be targeted to recipient cells efficiently, protect cytokine cargos from degradation, and/or change signaling downstream of cytokine receptors, EVs may serve to enhance or amplify cytokine signaling in tissues. As cytokines mediate immune cell communication during virtually all inflammation-mediated pathologies, understanding the mechanisms by which EVs control cytokine function could lead to insights into how to treat both allergic lung and other inflammatory airway diseases.

## Funding sources

This research was supported by NIH DP2 HL152426 (HHP), NIH T32GM008554 (KEB), and institutional funds from the Department of Pathology, Microbiology and Immunology at Vanderbilt University Medical Center (HHP).

## Acknowledgements

We thank Dr. Evan Krystofiak and the Vanderbilt Cell Imaging Shared Resource (CISR) core for electron microscopy imaging. We thank Dr. Andries Zijlstra and Dr. Paula Hurley for use of the CellStream for single vesicle flow cytometry. We thank Dr. Mary Philip for providing *Lck-Cre* mice. We thank Ms. Cherie Saffold for RNA extraction. We thank Dr. Neil Sprenkle for help with tissue collection. This work utilized equipment at the Vanderbilt Center for Extracellular Vesicle Research, the Vanderbilt Cell and Developmental Biology (CBD) Core, and the Vanderbilt University Medical Center Department of Pathology, Microbiology, and Immunology. We thank the Vanderbilt University Medical Center Animal Care and Use Program and veterinarians, veterinary technicians, and animal care technicians for mouse husbandry. Graphical figures were made using Biorender (www.Biorender.com).

## Contributions

KEB and HHP conceived and designed the research. KEB and BGG performed experiments. KEB and HHP analyzed and interpreted data. KEB and HHP prepared and drafted the manuscript. KEB, BGG, and HHP edited and revised the manuscript. All authors reviewed and approved the manuscript.

## Notes

**Disclosure Statement:** The authors have no conflicts of interest to disclose.

### Competing Interest Statement

The authors have declared no competing interest.

