## Supplemental Materials for "Th2 cell extracellular vesicles promote eosinophil survival through the cytokine cargo IL-3 and prolong airway eosinophilia"

### Supplementary Methods

#### 1. Animals

All animal studies were performed after obtaining approval from the Vanderbilt Institutional Animal Care and Use Committee. *mTmG* (Gt(ROSA)26Sortm4(ACTB-tdTomato,-EGFP)Luo/J) mice (JAX 007576) were crossed with *Lck-iCre* (B6.Cg-Tg(Lck-icre)3779Nik/J) mice (JAX 012837) to generate mice on a C57BL/6 background in which T cells are specifically labeled with mGFP. All non *Lck-Cre mTmG* mice used were wildtype C57BL/6J (JAX 000664). Mice were used between 6 and 16 weeks of age and were housed in a specific pathogen free facility. Allergen challenge and sensitization and retro-orbital injection were performed under isoflurane anesthesia. All fluid and tissue collection was performed after mice were euthanized.

#### 2. Isolation of murine CD4+ T cells

Spleens and inguinal, cervical, and axillary lymph nodes were dissected from female C57BL/6 mice between the ages of 6 and 12 weeks, and a single cell suspension was prepared by pressing the spleens and lymph nodes through a 70 µm nylon cell strainer. CD4+ T cells were isolated from spleen and lymph node single cell suspensions by positive bead selection using the Dynabeads FlowComp Mouse CD4 Kit (ThermoFisher Scientific 11461D) per manufacturer instructions.

#### 3. Polarization and activation of Th2 cells

Non-tissue culture treated plates were prepared by incubating with 1 µg/ml goat anti-hamster IgG (ThermoFisher 31115) overnight at 4C, washing, and then incubating with 1 µg/ml anti-CD3 (BioXCell Clone 145-2C11) and 1 µg/ml anti-CD28 (BioXCell Clone 37.51) at 37 oC for 2 hours. Isolated CD4+ T cells were stimulated with plate-bound anti-CD3 and anti-CD28, 10 ng/ml IL-4 (Peprotech 214-14), and 10 ng/ml anti-interferon gamma (BioXCell Clone XMG1.2) in opTmizer serum-free, phenol-free media with added expansion supplement (ThermoFisher A3705001) for 3 days. On day 3, cells were removed from anti-CD3/anti-CD28 stimulation and rested in opTmizer media with 10 ng/ml IL-4 and 60 U/ml IL-2 (National Cancer Institute at Frederick) for an additional day. Media was changed and replaced with fresh opTmizer media with 10 ng/ml IL-4 and 60 U/ml IL-2 on day 4 and cells were rested for an additional day. On day 5 of culture, polarized Th2 cells were placed back onto anti-CD3/anti-CD28 plates in opTmizer media with 60 U/ml IL-2 and re-stimulated for 6 hours.

#### 4. Isolation of EVs from murine rested and activated Th2 cell cultures

On day 5 of culture before restimulation, Th2 cells were centrifuged at 250xg for 10 minutes to pellet cells, and supernatant was collected. This supernatant was subjected to centrifugation at 2,000xg for 30 minutes to pellet large cellular debris and was used for isolation of rested Th2 cell EVs. Th2 cells resulting from 250xg centrifugation were re-stimulated for 6 hours, and media was subsequently collected and subjected to 250xg and 2,000xg differential centrifugation and was used for isolation of activated Th2 cell EVs. EVs were isolated from cell culture supernatant by size exclusion chromatography using the Izon 35 nm qEV100 column (IC100-35) or qEV10 column (IC10-35) per manufacturer instructions. The first 3x50 ml fractions following the void volume were collected for the qEV100 column, and the 2nd, 3rd, and 4th 5 ml fractions after the void volume were collected for the qEV10 column, as instructed by the manufacturer. Fractions were then concentrated using ultrafiltration with a 100 kDa filter (Millipore Sigma UFC710008).

**5. T cell viability assay**

Rested and activated Th2 cell viability was assayed using the LIVE/DEAD cell imaging kit 488/570 (Invitrogen R37601) per manufacturer instructions. After staining cells were imaged with the EVOS FL fluorescent microscope (ThermoFisher). The number of red and green cells in the images were quantified using Fiji (https://imagej.net/software/fiji/).

**6. Nanoparticle tracking analysis**

The concentration of EVs was analyzed by nanoparticle tracking analysis using a ZetaView (ParticleMetrix) with a 405 nm laser in scattering mode at RT. EVs were diluted 1000 to 10,000 times in PBS and measurements were performed using 11 camera positions with a sensitivity setting of 65. Data were analyzed using ParticleMetrix's NTA software.

**7. Transmission Electron Microscopy**

Activated and rested Th2 cell culture media was centrifuged at 250xg for 10 minutes and 2000xg for 30 minutes and then fractionated using Izon qEV100/35 nm Sepharose size exclusion columns (Izon IC100-35) per manufacturer instructions. The first 3 fractions were pooled and concentrated using ultrafiltration with 100 kDa pore size (Millipore Sigma UFC710008). Resulting concentrates were layered into an iodixanol gradient consisting of 40%, 20%, 10%, and 5% OptiPrep (StemCell Technologies 07820) diluted with 0.25 M sucrose, 50 mM EDTA, 10 mM Tris HCl pH 7.5 in PBS. Samples were loaded on the bottoms of the density gradients. Gradients were ultracentrifuged at 100,000xg for 18 hours. Gradient layers with buoyant density of EVs were isolated, diluted in PBS, and ultracentrifuged at 100,000xg for 4 hours. Resulting pellets were resuspended in PBS. EVs were adhered to freshly glow discharged, 300 mesh carbon coated grids for 10 seconds followed by 2 brief washes in $ddH_2O$ and stained with 2% uranylacetate. TEM was performed with the Tecnai T12 microscope, which operates at 100 keV using an AMT nanosprint CMOS camera.

**8. Induction of allergic airway inflammation with ovalbumin sensitization and challenge**

A solution of 10 mg/ml ovalbumin protein (OVA) from chicken egg white (Millipore Sigma A5503) was prepared by resuspending 50 mg OVA in 5 ml pharmaceutical-grade PBS (VWR K812). For sensitization, 5 µl of 10 mg/ml OVA was combined with 95 µl pharmaceutical-grade PBS and 100 µl Imject Alum (ThermoFisher 77161) per mouse (50 µg OVA per mouse), and this solution was mixed by gentle rocking at RT for at least 30 minutes prior to injection. Male and female C57BL/6 *Lck-Cre mTmG* mice between the ages of 6 and 12 weeks were injected intraperitoneally with 200 µl OVA+alum solution. For challenge, 5 µl of 10 mg/ml OVA was combined with 15 µl pharmaceutical-grade PBS per mouse (50 µg OVA per mouse). Mice were challenged by oropharyngeal aspiration of 20 µl OVA for allergic airway mice or 20 µl PBS vehicle control on days 7, 8, and 9 following sensitization. Mice were euthanized on day 10 following sensitization.

**9. Induction of allergic airway inflammation with house dust mite challenge**

A solution of 100 mg/ml house dust mite extract (HDM) from *Dermatophagoides Pteronyssinus* (Greer Laboratories Inc. XPB82D3A2.5 Lot 348717, Derp1 1449 mcg/vial, Dry weight 159.7

mg/vial, protein bradford 29 mg/vial, endotoxin 11400 EU/vial) was prepared by resuspending 159.7 mg HDM in 1.597 ml pharmaceutical-grade PBS (VWR K812). Single-use working stocks of 10 mg/ml HDM were prepared by diluting 100 mg/ml HDM in pharmaceutical-grade PBS. For challenge, 4 ul 10 mg/ml HDM was combined with 16 µl pharmaceutical-grade PBS per mouse (40 µg per mouse). Male and female C57BL/6 *Lck-Cre mTmG* mice between the ages of 6 and 12 weeks were challenged by oropharyngeal aspiration of 20 µl HDM for allergic airway mice or 20 µl PBS vehicle control on days 0, 2, 4, 7, 9, 11, 14, 15, and 16. Mice were euthanized on day 17.

**10. Induction of type 2 airway inflammation with papain challenge**

A solution of 5 mg/ml papain from papaya latex (Millipore Sigma P4762) was prepared by resuspending 50 mg papain in 10 ml pharmaceutical-grade PBS. For challenge, 3.5 µg papain in 40 µl pharmaceutical-grade PBS was administered by oropharyngeal aspiration to wildtype C57BL/6 mice between the ages of 6 and 12 weeks. Mice were euthanized 24 to 72 hours following challenge.

**11. Collection of bronchoalveolar lavage fluid cells and EVs**

Mice were euthanized by $CO_2$ inhalation. The diaphragm was punctured with a 23G needle, and a 0.5 cm incision was made in the trachea. A 23G catheter was inserted into the incision in the trachea, and 1 ml 0.22 µm filtered ice-cold PBS was flushed in and out of the airways 4 times to collect a total of ~3.5 ml bronchoalveolar lavage fluid (BALF). BALF was centrifuged at 250xg for 10 minutes to pellet cells, which were used in subsequent flow cytometry assays. The resulting supernatant was centrifuged at 2,000xg for 30 minutes to pellet large cellular debris, and the resulting supernatant was used in EV studies.

**12. Collection of lung tissue for flow cytometry**

To distinguish circulating immune cells from tissue-resident immune cells in the lung, circulating immune cells were labeled with anti-CD45 PE (Tonbo Biosciences, clone 30-F11, 50-0451) via retro-orbital injection of anti-CD45 PE. Mice were anesthetized and injected with 150 µl of anti-CD45 PE diluted 1:6 in pharmaceutical grade PBS using a 30G insulin syringe into the retro-orbital sinus. Mice were allowed to recover for 5 minutes and were then euthanized by $CO_2$ inhalation. Following BALF collection, the lungs and heart were removed *en bloc* and placed into complete media on ice. The right lobe was dissected away from the heart and left lobe, minced using a razor blade, and digested by rocking at 37C in complete media with 0.5 mg/ml collagenase from *Clostridium histolyticum* (Sigma C2674) and 5 U/ml DNase I (Qiagen 79254) for 45 minutes. A single cells suspension was then prepared by pressing the digested tissue through a 100 µm nylon cell strainer.

**13. Single Vesicle Flow Cytometry**

Single vesicle flow cytometry was performed using a CellStream (Luminex) flow cytometer with 456/51, 528/46, 583/24, 611/31, 792/87, and 773/56 lasers. Calibration was performed using CellStream Calibration Reagent (Luminex CS-400104) per manufacturer instructions prior to every experiment. Samples were acquired using small particle detection mode with FSC and SSC laser intensities set to 1% and all other laser intensities set to 100%. Samples were acquired for 2 minutes.

**14. Cell Flow Cytometry**

Cells isolated from BALF or lung tissue were counted using a hemocytometer, divided, and then stained with two panels of antibodies: (1) Lymphoid panel – anti-CD4 FITC (eBioscience, clone RM4-5, 35-0042), anti-CD8 PE-Cy7 (Invitrogen, clone 53-6.7, 25-0081-82), and anti-CD45 APC (Invitrogen, clone 30-F11, 17-0451-82) (2) Myeloid panel – anti-F4/80 PE-Cy7 (Tonbo Biosciences, clone BM8.1, 60-4801-4100), anti-Siglec F PerCP-Cy5.5 (BD Pharmingen, clone E50-2440, 565526), anti-CD11b Pacific Blue (Tonbo Biosciences, clone M1170, 75-0112-4100), anti-CD11c FITC (eBioscience, clone N418, 11-0114-85), and anti-CD45 APC (Invitrogen, clone 30-F11, 17-0451-82). All antibodies were diluted 1:8. Both panels also included the viability dye eFluor 780 (eBioscience 65-0865) at a dilution of 1:500 to distinguish between live and dead cells. Compensation controls were prepared from lymphocytes collected from mouse lymph nodes at the time of the experiments. Flow cytometry was performed using BD Biosciences FACSCanto II flow cytometer with three excitation lasers (488 nm, 633 nm, and 405 nm) and 10 fluorescence channels. Flow cytometry data were analyzed using FlowJo software (FlowJo LLC, Ashland, OR).

**15. Bone marrow-derived eosinophil (BMDE) differentiation**

Femurs and tibias were dissected from C57BL/6 mice between the ages of 6 and 12 weeks old. The bones were cut, and 10 ml Iscove's Modified Dulbecco's Medium (Gibco 12440053) was flushed through the bone cavity to dislodge the bone marrow. The bone marrow was pressed through a 100 µm nylon cell strainer to create a single cell suspension. Red blood cells were lysed by incubating the cells in Ack lysing buffer (ThermoFisher A1049201), and cells were washed. Resulting bone marrow cells were incubated in RPMI 1640 (ThermoFisher 11875119) supplemented with penicillin/streptomycin, HEPES (ThermoFisher 15630080), sodium pyruvate, L-glutamine, betamercaptoethanol, non-essential amino acids (ThermoFisher 11140050), and 20% FBS (Gibco). Cells were cultured with 100 ng/ml Flt3-ligand (Peprotech 250-31L) and 100 ng/ml stem cell factor (Peprotech 250-03) for four days and then switched to media containing 10 ng/ml IL-5 (Peprotech 215-15) for 8 days with media change every other day. Eosinophils were used in experiments on day 12 of culture.

**16. Eosinophil functional studies**

***16a. Microscopy***

Day 12 BMDEs were deposited onto a glass slide by cytocentrifugation at 800 RPM for 2 minutes. Slides were stained using HARLECO hemacolor stain set (Millipore Sigma 65044) per manufacturer instructions.

***16b Chemotaxis assay***

500 µl BMDE media was added to wells of a 24-well 0.4 µm pore size transwell plate (ThermoFisher 140620) and the transwell inserts were inserted into the wells and allowed to equilibrate at 37C for 2 hours. Recombinant murine CCL11 (Peprotech 250) was loaded into the bottom or top well at concentrations of 0, 20, or 200 ng/ml. 250,000 day 12 BMDEs were loaded into the top well and incubated overnight at 37C. Cell migration was quantified by flow cytometry using the Luminex CellStream.

***16c. Eosinophil peroxidase assay***

Day 12 BMDEs were washed and resuspended in phenol-free RPMI with L-glutamine, 25 mM HEPES, and 0.1% BSA at a concentration of 500,000 cells/ml. Fifty µl cells were added in triplicate to the wells of a non-tissue culture treated 96-well plate, and the plate was incubated at 37C for 1 hour. 50 µl calcium ionophore A23187 (Millipore Sigma C7522) was added to the wells along a dose titration, and BMDEs were incubated at 37C for 30 minutes. The peroxidase substrate was prepared by adding o-Phenylenediamine (OPD) (Millipore Sigma P9029) to 400 mM Tris-HCl, pH 8.0 (Corning 46-031-CM) in distilled water such that the final concentration of OPD was 400 µM. Right before use, $H_2O_2$ was added for a final concentration of 1.1 µM. 100 µl peroxidase substrate was added to the BMDEs and incubated for 4 minutes. 100 µl $H_2SO_4$ was added to stop the reaction, and absorbance was measured using a BioTek Synergy HTX microplate reader at 490 nm. Total eosinophil peroxidase was measured by lysing the cells with 0.2% Triton X-100 prior to addition of peroxidase substrate.

**17. EV treatment of bone marrow-derived eosinophils**

EVs were purified from rested and activated Th2 cell culture by SEC and ultrafiltration as described previously. Samples were sterile 0.22 µm filtered. EV concentration was measured using nanoparticle tracking analysis. On day 12 of culture, 200,000 BMDEs were plated in 200 µl complete RPMI in 96-well plates and were treated with $5x10^9$ EVs/ml for 6 or 24 hours.

**18. Administration of EVs *in vivo***

EVs were purified from activated Th2 cell culture by SEC and ultrafiltration as described previously. Media not conditioned with cells was subjected to parallel processing as a control. Samples were sterile 0.22 µm filtered. EV concentration was measured using NTA, and the concentration of EVs in the sample was adjusted to $2x10^9$ EVs/40 µl using pharmaceutical grade PBS. Mice were anesthetized and administered 40 µl EVs or media control by oropharyngeal aspiration.

**18. Eosinophil death assays**

***18a. Eosinophil membrane permeability assay***

EVs were purified from rested and activated Th2 cell culture media as described previously. BMDEs on day 12 of culture were treated with $5x10^9$ EVs/ml for 24 hours. Following treatment, cells were washed and stained with Fixable Viability Dye eFluor 780 (eBioscience 65-0865) at a 1:500 dilution per manufacturer instructions. Cell death as measured by membrane permeability to the dye was assayed by flow cytometry.

***18b. Eosinophil caspase 3 activity assay***

EVs were purified from rested and activated Th2 cell culture media as described previously. BMDEs on day 12 of culture were treated with $5x10^9$ EVs/ml for 6 hours. Following treatment, caspase 3 activity was assayed using the EnzChek Caspase 3 Assay Kit (Invitrogen E13183) per manufacturer instructions. Caspase 3 activity as measured by substrate cleavage by caspase 3 was assayed by flow cytometry.

***18c Eosinophil Annexin V binding assay***

EVs were purified from rested and activated Th2 cell culture media as described previously. BMDEs on day 12 of culture were treated with $5x10^9$ EVs/ml for 6 hours. Following treatment, the presence of phosphatidyl serine on eosinophil membranes was measured via Annexin V binding using the Dead Cell Apoptosis Kit Annexin V FITC for flow cytometry (Invitrogen V13242). The cells were also stained with Fixable Viability Dye eFluor 780 as described previously, and apoptotic cells were defined as cells positive for Annexin V and negative for eFluor 780 by flow cytometry.

**19. Detergent treatment and proteinase K treatment of EVs**

For Fig. 1B and C and Supplemental Fig. 1C, cell culture media or BALF was centrifuged at 250xg and then 2000xg and subsequently treated with either Triton X-100 detergent (Sigma T8787) or proteinase K (Fisher Bioreagents BP-1700-100, lot #230139) at varying dilutions and mixed thoroughly. Proteinase K-treated EVs were incubated at 37C for 1 hour. Treated samples were then subjected to single vesicle flow cytometry.

For Fig. 5E and Supplemental Fig. 6E-G, EVs purified from Th2 cell culture by SEC as described above were incubated with 0, 2, or 10 µg/ml PK at 37C for 1 hour. Non-conditioned cell culture media processed parallelly with EV-containing cell culture media was also treated with PK and used as a sham control. Following the incubation, EVs were loaded into a 100 kDa pore size centrifugal filter (Amicon UFC5100) and centrifuged at 14,000xg for 15 minutes. The flow-through containing PK was discarded, and the retentate containing EVs was used for experiments.

**20. Ruxolitinib treatments**

For ruxolitinib dose titration, ruxolitinib at 10 mM in DMSO (APExBIO A3012) and DMSO as a vehicle control were diluted in sterile PBS to form a 5-fold dilution series. Day 12 BMDEs were treated with 10 ng/ml IL-5 and either ruxolitinib or DMSO and incubated at 37C for 24 hours after which viability was assayed by membrane permeability assay. For ruxolitinib and EV co-treatment, day 12 BMDEs were treated with 1.6 µM ruxolitinib or DMSO for 30 minutes at 37C. BMDEs were then incubated with $5 \times 10^9$ rested or activated Th2 cell EVs/ml purified by SEC or 10 ng/ml IL-5 for 24 hours at 37C after which viability was assayed by membrane permeability assay.

**21. EV IL-3 Enzyme Linked Immunosorbent Assay (ELISA)**

EVs were purified from activated or rested Th2 cell culture media by SEC. For Fig. 5D and Supplemental Fig. 6C, Izon 35 nm qEV10 columns were used to fractionate media, and fractions 1-9 were collected and concentrated using 15 ml 100 kDa ultrafilters (Millipore Sigma UFC9100). For Fig. 5C, F, and G, Izon 35 nm qEV100 columns were used to fractionate media, and fractions 1-3 were collected, concentrated using 70 ml 100 kDa ultrafilters (Millipore Sigma UFC710008), and pooled. The mouse IL-3 Quantikine ELISA kit (R&D Systems M3000) was used to detect IL-3 in EV samples. ELISA plates were prepared per manufacturer instructions, and concentrated EV samples were added to the plate in duplicate. IL-3 detection was performed per manufacturer instructions, and absorbance was measured using a BioTek Synergy HTX microplate reader at 450 nm and 540 nm wavelengths. A standard curve was prepared per manufacturer instructions and used to determine concentrations for absorbance values. Absorbance values at 540 nm were subtracted from values at 450 nm per manufacturer instructions.

**22. IL-3 Blocking**

EVs purified from rested and activated Th2 cell culture media by SEC were incubated with 1, 10, or 100 µg/ml anti-mouse IL-3 (BioXcell BE0282, clone MP2-8F8) or isotype control (BioXcell BE0083, clone MOPC-21) for 30 minutes at 37C. Sterile PBS was also incubated with anti-IL-3 or isotype control for use as a vehicle control. EVs at a concentration of $5x10^9$ EVs/ml or equal volume of vehicle were then administered to day 12 BMDEs, and BMDEs were incubated for 24 hours at 37C. Viability was then assessed using membrane permeability assay.

**23. IL-3 Potency**

EVs were purified from activated Th2 cell culture using SEC, and the concentration of IL-3 associated with EVs was measured by ELISA as outlined above. BMDEs were treated with increasing concentrations of EV-associated IL-3 or recombinant murine IL-3 (Peprotech 213-13) and incubated for 24 hours at 37C. Viability was assessed using membrane permeability assay.

**24. Bulk RNA-sequencing**

Day 12 BMDE treated with either EVs purified from activated Th2 cell culture or parallely processed cell culture media for 24 hours were homogenized with TRIzol Reagent (Invitrogen 15596026). Total RNA was isolated using the RNeasy Mini Kit (Qiagen 74104) and submitted to the Vanderbilt Technologies for Advanced Genomics Core for sequencing on the NovaSeq 6000 (Illumina). Data were processed on the Vanderbilt computing cluster (ACCRE) using the lab's RNA-seq pipeline (https://github.com/parkdj1/RNASeq-Pipeline). Briefly, FastQC was used to evaluate data quality, adapters were trimmed using Cutadapt, and alignment with quantification was performed using Salmon. Primary data was deposited in the Gene Expression Omnibus (https://www.ncbi.nlm.nih.gov/geo/). Raw and processed data can be accessed using the accession number GSE253180 with the reviewer token ojexoeygplqbrgv. Downstream analysis included differential expression using the R package DESeq2 (https://www.r-project.org/) and gene set enrichment (www.broadinstitute.org/gsea). Heatmaps were made using the R package pheatmap, and volcano plots were made using GraphPad Prism 10 software (GraphPad Software, La Jolla, CA).

**25. Statistical analyses**

Data were expressed as mean values plus or minus standard errors of the mean or as median and interquartile range depending on the normality of the data. Statistical significance was determined through tests denoted in the figure legends performed by GraphPad Prism 10 software (GraphPad Software, La Jolla, CA). P-value of $< 0.05$ was considered statistically significant.

**Supplemental Figure 1.**

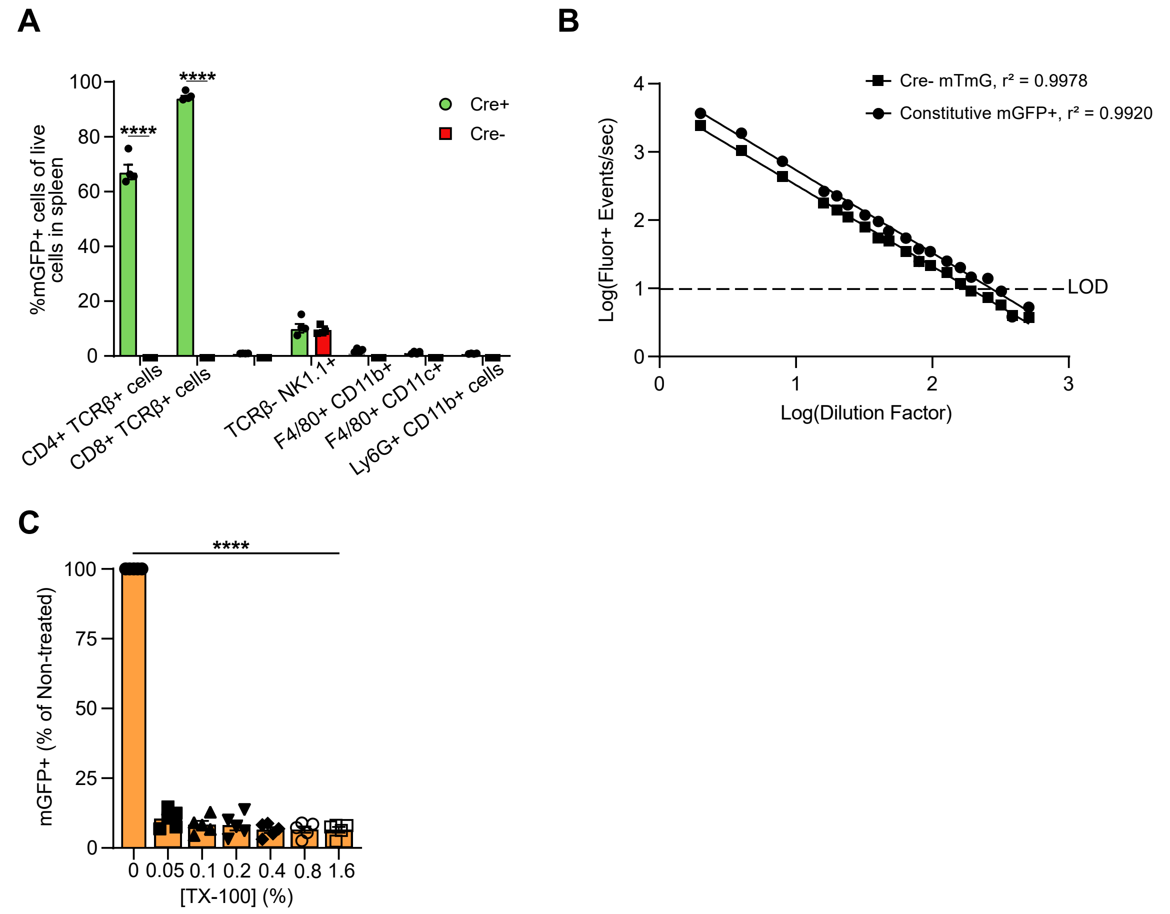


**Supplemental Figure 1. BALF mGFP$^{+}$ and mTomato$^{+}$ EVs are detected in the linear range of single vesicle flow cytometry and are sensitive to Triton X-100 lysis. (A)** Flow cytometric analysis of mGFP labeling of immune cell types in the spleens of *Lck-Cre mTmG* mice, including CD4$^{+}$ T cells (CD4$^{+}$ TCRβ$^{+}$), CD8$^{+}$ T cells (CD8$^{+}$ TCRβ$^{+}$), B cells (CD19$^{+}$ TCRβ$^{-}$ ), natural killer cells (TCRβ$^{-}$ NK1.1$^{+}$ ), macrophages/monocytes (F480$^{+}$ CD11b$^{+}$ /CD11c$^{+}$ ), and neutrophils (Ly6G$^{+}$ CD11b$^{+}$ ). n = 4, 2-way ANOVA with Tukey's test for multiple comparisons. **(B)** Flow cytometric analysis of mGFP$^{+}$ and mTomato$^{+}$ event detection per second in BALF collected from a *Cre$^{-}$ mTmG$^{+}$* mouse, in which all EVs are mTomato$^{+}$, and a constitutive mGFP expressing mouse, in which all EVs are mGFP$^{+}$ over a dilution series. n = 1, simple linear regression. **(C)** Quantification of mGFP$^{+}$ particles detected in OVA *Lck-Cre mTmG* BALF following treatment with varying concentrations of TX-100, n = 5, one-way ANOVA (vs. 100%) with Bonferonni correction for multiple testing, error = SEM. **** $p < 0.0001$.

**Supplemental Figure 2.**

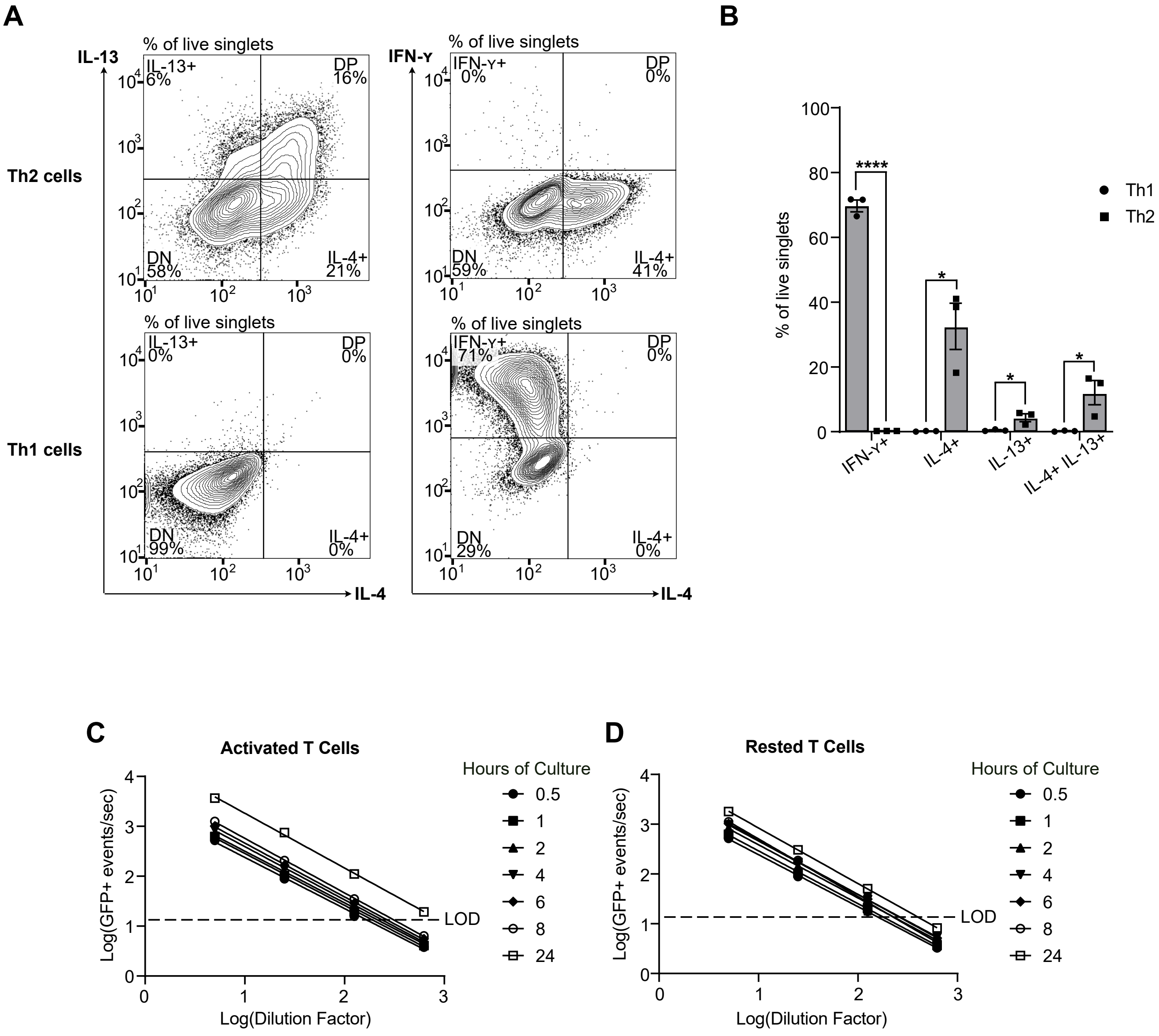


**Supplemental Figure 2. T cells activate and polarize to Th2 cells in serum-free, chemically defined media. (A)** Representative flow cytometry plots of IFNγ, IL-4, and IL-13 production by primary mouse CD4⁺ T cells isolated from spleens and lymph nodes and then polarized with T cell receptor stimulation and either IL-4 and anti-IFNγ for Th2 cells (top) or IL-12 and anti-IL-4 for Th1 cells (bottom). **(B)** Quantification of IFNγ, IL-4, and IL-13 producing T cells following flow cytometric analysis. n = 3, 2-way ANOVA with Tukey's test for multiple comparisons. **(C, D)** Flow cytometric analysis of mGFP event detection per second in media collected from **(C)** activated and **(D)** rested *Lck-Cre mTmG* T cells over a dilution series that includes 5-, 25-, 125-, and 625-fold dilutions. n = 3, simple linear regression, $r^2$ all > 0.98. All errors bars represent SEM, * $p < 0.05$, **** $p < 0.0001$.

**Supplemental Figure 3.**

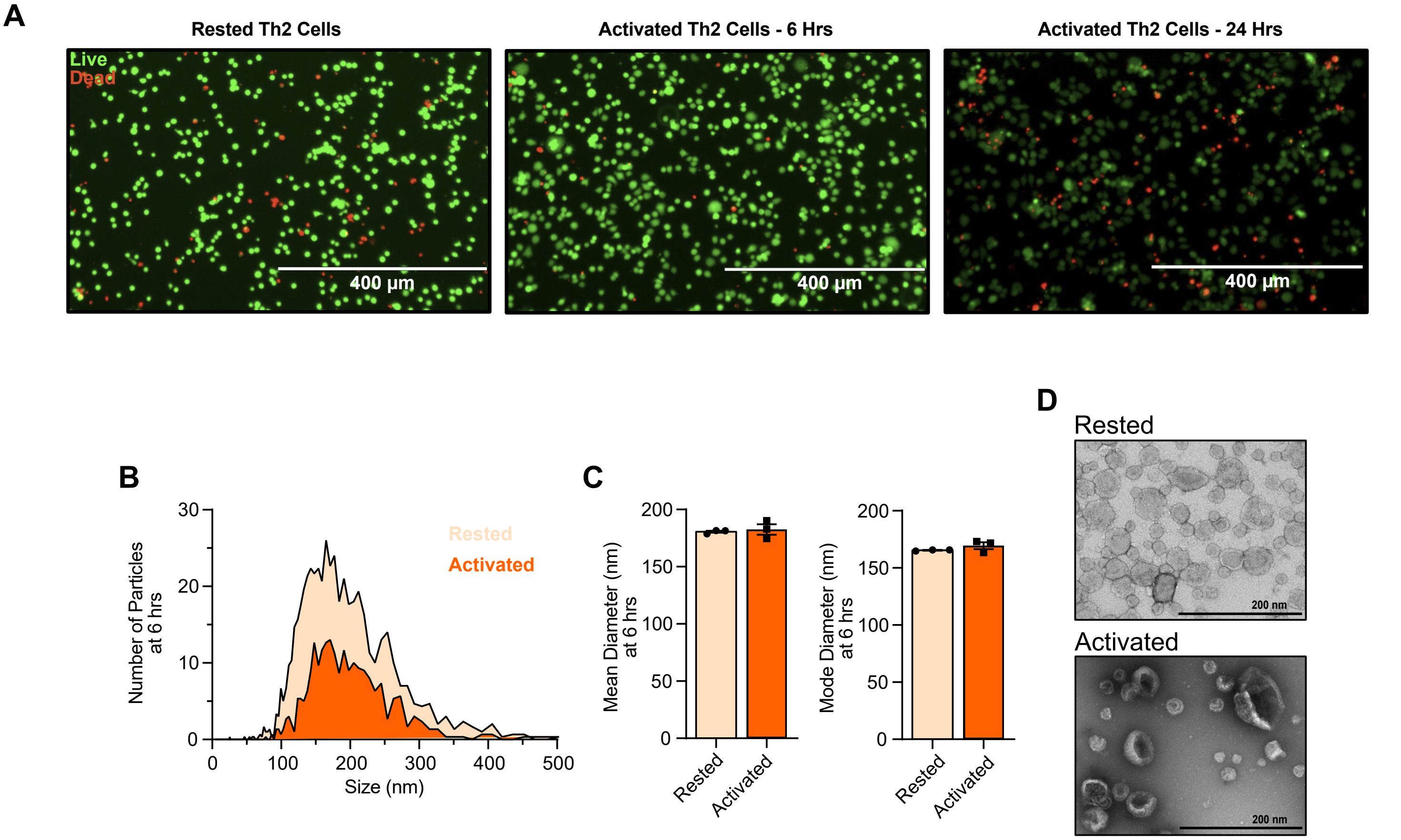


**Supplemental Figure 3. Activation state does not change gross morphological characteristics of Th2 cell EVs. (A)** Representative images of live/dead-stained rested (left), activated for 6 hours (middle), and activated for 24 hours (right) Th2 cells. **(B)** Average size distribution of EVs produced by rested (light orange) and activated (dark orange) Th2 cells at 6 hrs of culture as measured by nanoparticle tracking analysis. Histograms represent averages of 3 samples. **(C)** Mean and mode size of EVs produced by rested and activated Th2 cells at 6 hrs of culture as measured by nanoparticle tracking analysis. n =3, two-tailed t test. **(D)** Transmission electron micrographs of EVs purified by density gradient from rested (top) and activated (bottom) Th2 cell culture media, magnification 1100x. All error bars represent SEM.

### Supplemental Figure 4.

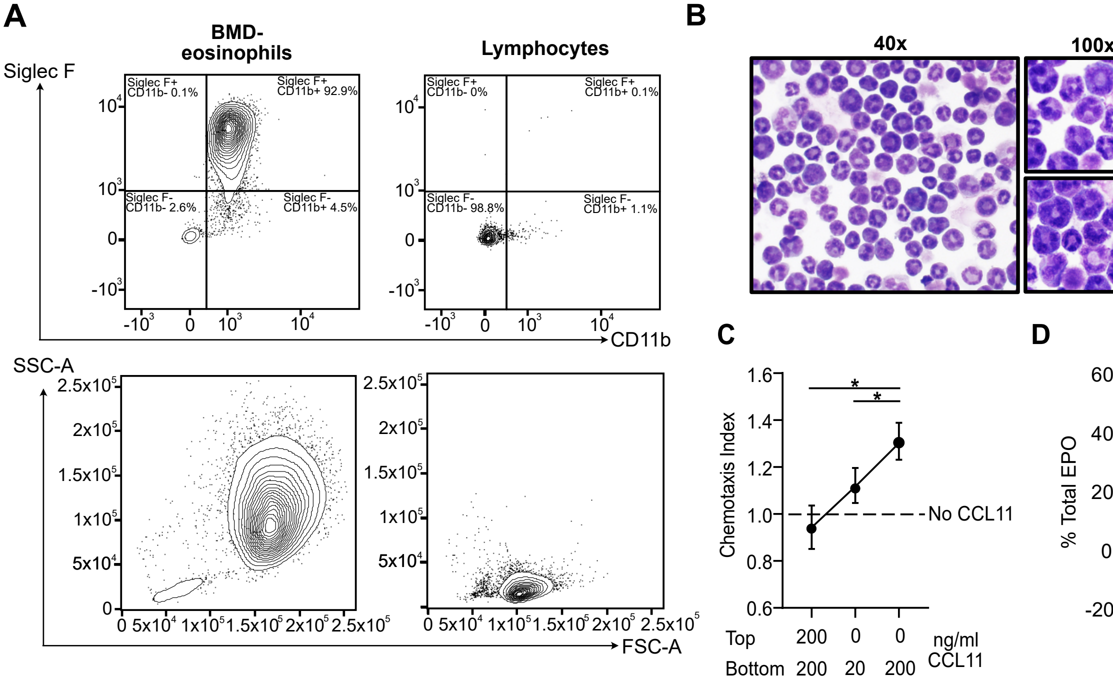


**E**

Estimated physiological concentration of T cell EVs in the airway lining fluid during allergic inflammation = 4.62x10⁶ T cell EVs/airway (mean # mGFP⁺ EVs in OVA BALF) / 1 ml (volume of mouse lung) x 100 (dilution factor of BALF) x 5 (correction for sensitivity of EV flow cytometry vs. NTA)

**Supplemental Figure 4. Eosinophils differentiated from mouse bone marrow recapitulate eosinophil functions *in vitro*. (A)** Representative flow cytometry plots of day 12 bone marrow-derived eosinophils (left) and lymphocytes (right) stained for the eosinophil markers Siglec F and CD11b (top) and assayed for forward- and side-scatter (bottom). **(B)** Light micrographs of day 12 bone marrow-derived eosinophils stained with hemacolor staining kit at 40x and 100x magnification. **(C)** Quantification of day 12 bone marrow-derived eosinophil chemotaxis to the eotaxin CCL11 by a transwell migration assay. n = 3, one-way ANOVA with Tukey's multiple comparisons test. **(D)** Quantification of day 12 bone marrow-derived eosinophil eosinophil peroxidase (EPO) production in response to increasing doses of the calcium ionophore A23187, n = 3, log[agonist] vs response with variable response (4 parameters). **(E)** Formula for calculating the estimated physiological concentration of T cell EVs in airway lining fluid during allergic inflammation. All error bars represent SEM, * $p < 0.05$.

**Supplemental Figure 5.**

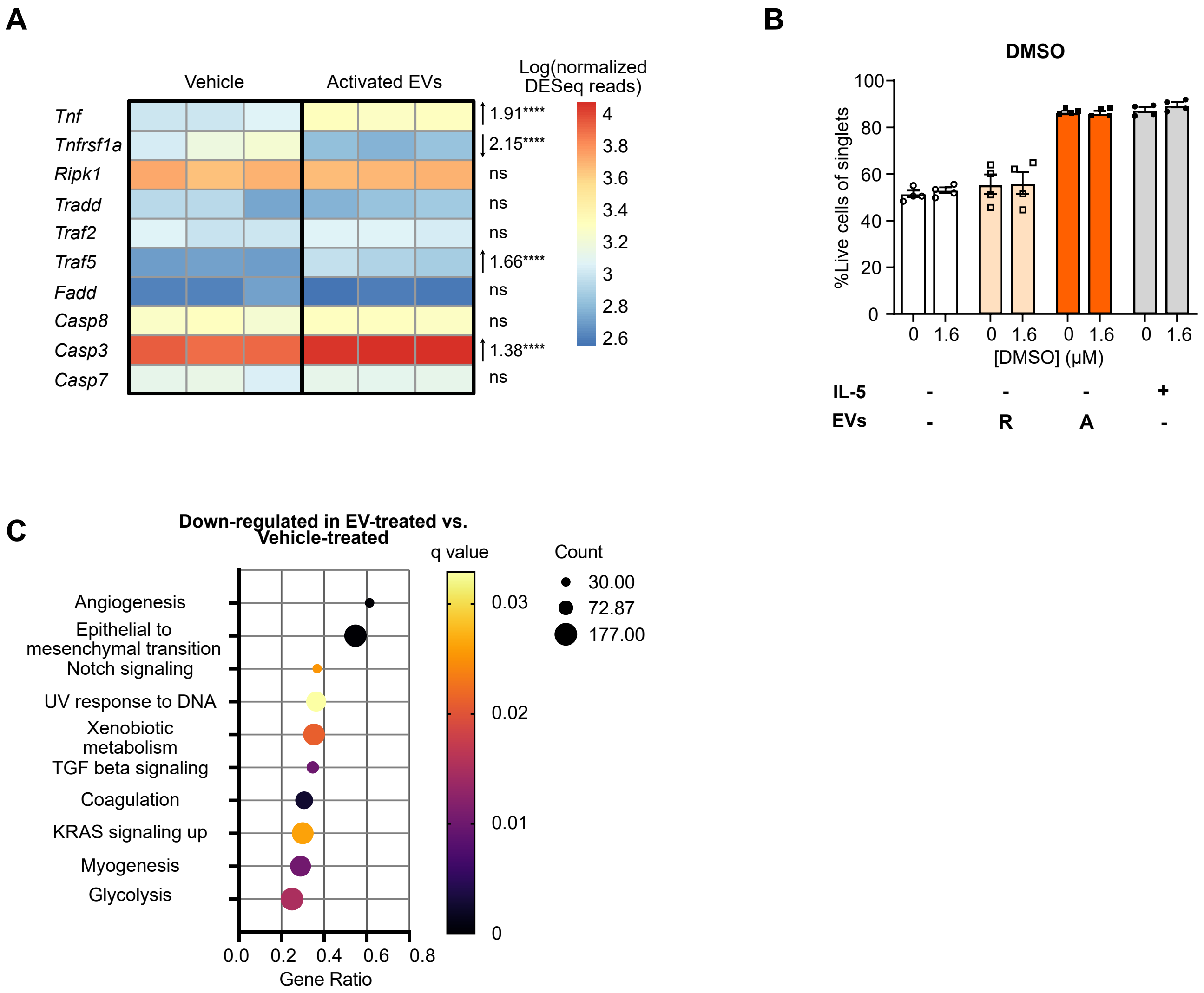


**Supplemental Figure 5. Activated T cell EVs regulate metabolism pathways and extrinsic apoptosis genes in eosinophils *in vitro*. (A)** Heatmap depicting Kegg extrinsic apoptosis pathway genes and corresponding normalized DESeq reads for those genes in vehicle treated eosinophils and Th2 cell EV treated eosinophils. **(B)** Quantification of eosinophil survival following 24 hrs treatment with rested or activated Th2 cell EVs or IL-5 and 1.6 μM DMSO, n = 4, 2-way ANOVA with Sidak's correction for multiple comparisons. **(C)** Gene set enrichment analysis of Th2 cell EV treated eosinophils vs vehicle treated eosinophils. Shown are gene sets downregulated in EV treated vs vehicle treated bone marrow-derived eosinophils with q-value ≤0.05. Differential gene expression analysis was performed using the R package DESeq2. All error bars represent SEM, **** $p < 0.0001$.

**Supplemental Figure 6.**

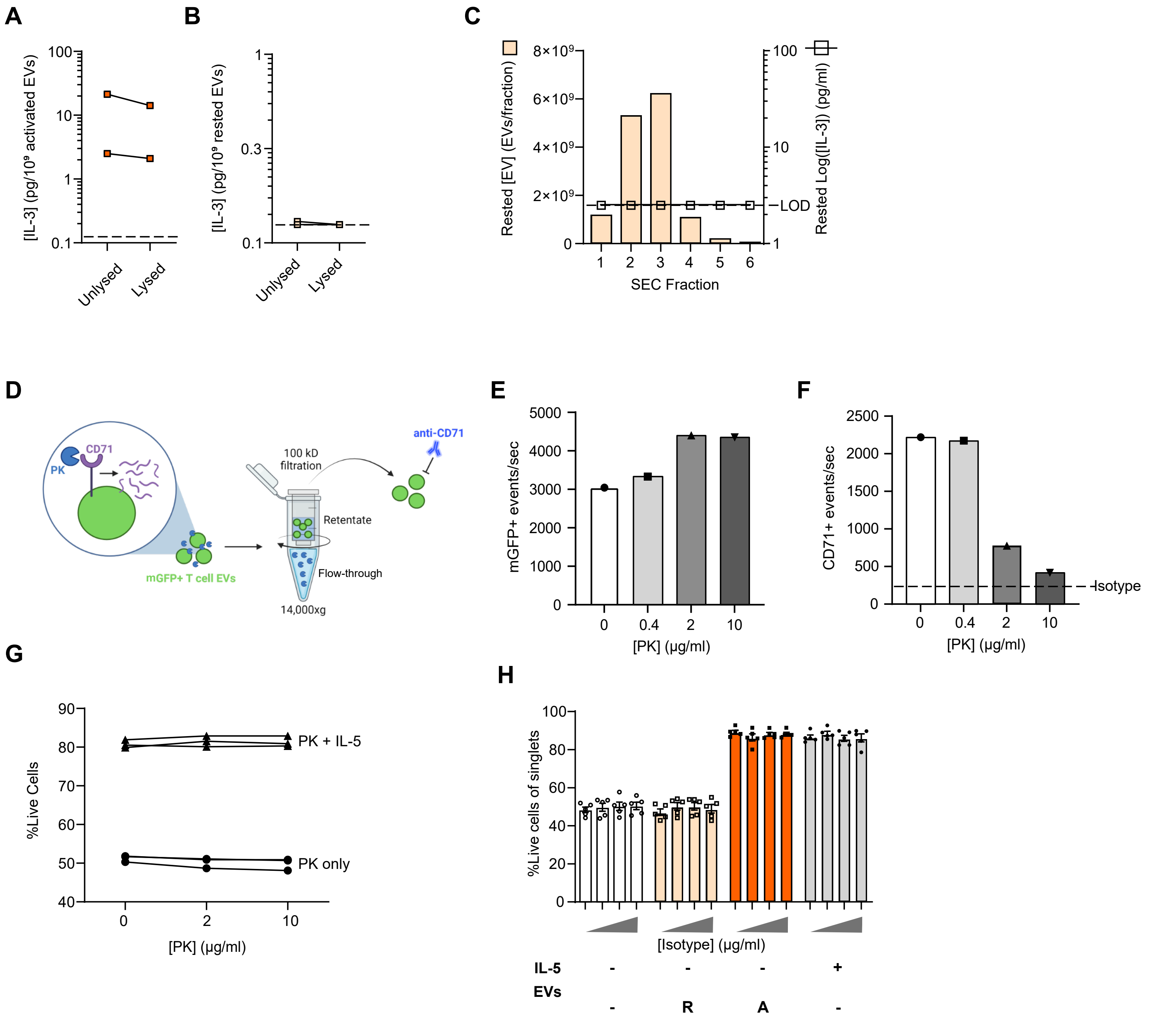


**Supplemental Figure 6. IL-3 is on the surfaces of EVs secreted by activated Th2 cells. (A, B)** Quantification of EV-associated IL-3 by ELISA with and without lysis of activated **(A)** and rested **(B)** Th2 cell EVs purified from cell culture media by SEC and ultrafiltration. **(C)** Quantification of EV number by NTA (left y-axis) and IL-3 concentration by ELISA (right y-axis) in SEC fractions 1-6 collected from rested Th2 cell culture media. **(D)** Primary eosinophil viability is sensitive to proteinase K (PK) treatment; therefore, EVs must be separated from PK before they are added to cell culture. Graphical depiction of EV PK treatment strategy. mGFP$^+$ T cell EVs were incubated with PK, after which they were subjected to centrifugal filtration using a 100 kD pore size. The flow-through containing PK was discarded, and the retentate containing EVs was stained with a fluorescent antibody against CD71, an abundant T cell EV surface cargo. EVs were then subjected to single vesicle flow cytometry to assess the integrity of the EVs and confirm surface protein cargo degradation. **(E)** Quantification of the number of mGFP$^+$ events detected per second in the retentate in *Lck-Cre mTmG* T cell culture media after treatment with increasing concentrations of PK. **(F)** Quantification of the number of CD71$^+$ events detected per second in the retentate in *Lck-Cre mTmG* T cell culture media after treatment with increasing concentrations of PK. For **(E)** and **(F)**, n = 1 set of T cell EVs pooled from 3 mice. The dotted line depicts fluorescent events detected per second with an isotype control antibody. **(G)** Quantification of cell death following eosinophil treatment with sham PK control retentate with or without IL-5. Non-conditioned cell culture media parallelly processed with EV-containing media was treated with PK as depicted in **(D)**. n = 3, 2-way ANOVA with Tukey's multiple comparison test (comparison of PK doses within PK only or PK + IL-5 conditions). **(H)** Quantification of eosinophil survival following 24 hrs of treatment with rested or activated EVs or vehicle (PBS) pre-incubated with increasing concentrations of isotype control antibody for anti-IL-3 (0, 1, 10, and 100 µg/mL). n = 5, 2-way ANOVA with Tukey's correction for multiple comparisons, 2 independent experiments.

**Supplemental Table 1. Results of differential gene expression analysis by DESeq2 in activated Th2 cell EV-treated vs. vehicle-treated eosinophils.**

**[Excel file]**

**Supplemental Table 2. Normalized read counts by DESeq2 in activated Th2 cell EV-treated and vehicle-treated eosinophils.**

**[Excel file]**
